# CNIH3 is a molecular signature of slow AMPA receptors

**DOI:** 10.64898/2026.07.22.739851

**Authors:** Marin Boutonnet, Jennifer D. Noonan, Niccolo P. Pampaloni, Andrew J.R. Plested

## Abstract

At excitatory synapses, glutamate activates receptors in the postsynaptic membrane including the AMPA receptor. Usually considered to be the fastest ion channel receptors in the brain, AMPA receptors can produce millisecond synaptic potentials that mimic the timing of spikes. However, we found abundant slow AMPA responses in CA1 pyramidal cells. These slow responses show a mosaic distribution in individual cells and even in single dendrites (Pampaloni et al., 2021). A survey of the literature reveals cryptic reports of similar slow responses in cerebellum, striatum and other brain regions (Pampaloni and Plested, 2022). These observations open a new perspective on the extent of AMPA receptor diversity. To enable a detailed interrogation of slow AMPA receptors in the brain, we sought to determine their molecular basis. We noted that slow AMPA is prevalent in ventral CA1 but rather sparse in dorsal hippocampus and absent in dentate gyrus granule cells. Checking published transcriptomic data, we noted that certain AMPA receptor auxiliary proteins also show gradients across these hippocampal regions. One understudied auxiliary protein identified from transcriptomics, CNIH3, produces uniquely slow AMPA responses in heterologous expression. Strikingly, we found that shRNAs against CNIH3 could ablate slow AMPA responses in ventral CA1, and over-expression of CNIH3 in dorsal CA1 introduced widespread mosaic slow AMPA responses. We conclude that CNIH3 is a sparsely-expressed marker of slow AMPA currents with the potential to convert individual synapses from fast followers into detonators that can spike the target cell, with implications for subcellular calculations, circuits, behaviour and brain pathologies.

## Introduction

Synaptic connections between neurons build up circuits in the brain but change their strength dynamically in time. The resulting “synaptic weights” define network activity and, ultimately, calculations and cognitive processes in the brain. The release of a synaptic vesicle triggered by a presynaptic action potential rapidly floods the synaptic cleft with neurotransmitter which is then cleared by diffusion and transport on the 100 µs timescale. The activation of cognate receptors can be just as quick, setting a lower limit on how fast a synapse can be (Taschenberger and von Gersdorff, 2000). But the magnitude and duration of the resulting postsynaptic signal is highly dependent on the properties of the postsynaptic receptors and their abundance (Henley and Wilkinson, 2016; Nusser, 2000; Coombs and Cull-Candy, 2021). At classical fast excitatory synapses populated by AMPA-type glutamate receptors, the characteristic decay time is less than 10 ms, but examples of sub millisecond synaptic currents are also reported (Geiger et al., 1997). These timescales relate well to the fast responses of AMPARs in heterologous expression (Mosbacher et al., 1994; Robert et al., 2001; Koike et al., 2000). The summation of these fast AMPAR responses at synapses permits the postsynaptic cells to translate action potential timing faithfully, essential for many common brain functions such as hearing (Hermann et al., 2007), reflexes (Li et al., 2003) and sensory information processing (Roth et al., 2020).

However, action potentials are much briefer than cognitive processes (which have a characteristic time of 200 ms and above (Buzsáki and Moser, 2013), and synchronous activity of neurons can be tuned at a low frequency, such as 4-8 Hz theta rhythms, underlying multiple cognitive functions. Many studies connect synapses to cognitive disorders, but how can synapses be coupled to cognition if their characteristic times are too short? Slow time constants in the brain might be generated by persistent activity and recurrent connectivity (related to “reservoir computing”) (Suárez et al., 2024), but this is implausible in many regions of the brain, because most neurons do not spike very much (around 1 Hz on average) and do not make enough recurrent connections (Seeman et al., 2018) (Peng et al., 2024). This paradox motivates the idea that other sources of slow time components are needed.

In addition to the widely expressed NMDA receptors, which operate over the 100 ms timescale, several further examples of slow glutamate receptors have been described, such as kainate receptors (limited to the mossy fiber-CA3 synapse), or delta subtype (very narrowly expressed and not activated by glutamate). We previously demonstrated that a new type of slow glutamate receptor, surprisingly variants of the AMPA receptor, is readily found within the hippocampus (Pampaloni et al., 2021). These slow AMPA receptors produce very long, non-desensitizing responses that accumulate, and synaptic decays lasting hundreds of milliseconds. These responses are reminiscent of the well-studied accumulating KAR current at the MF-CA3 hippocampal synapse (Castillo et al., 1997; Vignes and Collingridge, 1997), and similar to classical NMDA currents (for example GluN2A-containing, mature NMDA receptors, 100-200 ms (Gray et al., 2011). Because these slow AMPA responses allow exaggerated charge transfer, they can convert the synapses where they are expressed into conditional detonators, spiking the cell (Pampaloni et al., 2021), (Bischofberger et al., 2006). The slow AMPA responses recorded in the hippocampal pyramidal cells match well to cryptic reports of slow, non-NMDA currents that are seen in cerebellum, hippocampal interneurons, striatum, and so on (reviewed in (Pampaloni and Plested, 2022))

We previously reported a mosaic distribution of AMPA currents in CA1 pyramidal cells, ranging from typical fast responses, to hundreds of milliseconds decay responses in a single neuron. We also observed striking cell-to-cell variability, where only a minority of the recorded neurons displayed slow responses in response to a mild train of stimulation performed by glutamate uncaging (10 x 1 ms pulses at 10 Hz). There is mounting evidence that CA1 pyramidal neurons are a heterologous cellular population displaying highly distinct properties. Different gradients of genetic expression along the dorso-ventral ‘long axis’ are reported for instance (Cembrowski et al., 2016b). This heterogeneity of response kinetics would therefore potentially reflect synaptic properties in a subgroup of hippocampal pyramidal neurons supported by specific gene expression patterns.

AMPA receptors composed only of the principal GluA1-4 subunits cannot produce slow time constants of activation (but can provide short term depression from their relatively stable desensitization, 50-100 ms (Rothman et al., 2009)). However, co-expression with so-called auxiliary proteins (which include TARPs, cornichons and others (Kato et al., 2010; McGee et al., 2015; Carbone and Plested, 2016; Shi et al., 2010) can transform AMPA receptor activation. Depending on the particular auxiliary protein, properties such as resistance to desensitization, slower deactivation decays and altered ion permeation and block are seen. However few clear functional roles were demonstrated for these variant receptors at synapses. Likewise, the effects of TARPs on ion permeation and block were not clearly demonstrated at synapses. Instead the major role of TARPs appears to be as an essential anchor at the PSD (Bats et al., 2007; Chen et al., 2000). Some auxiliary proteins such as γ-4 and CNIH2 were demonstrated to produce slower synaptic decays under limited circumstances - for example CNIH2 slows the EPSCs of the Hilar Mossy cells (Boudkkazi et al., 2014; Milstein et al., 2007; Cho et al., 2007). In our previous work, expression of dominant-negative TARPs (designed to maintain receptor trafficking but not modulate gating) had only minor effects on slow AMPA currents (Pampaloni et al., 2021). Overexpression of γ-8, on the other hand, converted practically all uncaging sites to have a slow response. Given that this was on the background of existing γ-8 expression, and some other cells have expressed γ-8 (dentate gyrus granule cells) but completely lack slow AMPA responses, these data did not yield a clear interpretation. Therefore, to date the molecular identity of slow AMPA receptors, at least in hippocampal CA1 pyramidal cells, remains unclear.

Here we asked two main questions. First, is there a spatial or regional distribution of slow AMPA in the hippocampus? We found a strong gradient of slow AMPA responses in the mouse hippocampus, with slow responses very rare in dorsal CA1, but prevalent in ventral CA1. The same approach confirmed a clear absence of slow AMPA in DG. Building on this functional characterisation, we asked how this functional gradient is generated by the molecular composition of slow AMPA receptors. We probed public transcriptomics databases and identified that the Cornichon homolog-3 (CNIH3) is a candidate slow AMPA subunit, and confirmed it has the necessary properties in heterologous expression. Knockdown of CNIH3 in ventral hippocampus all but abolished the otherwise prevalent slow AMPA responses, whereas overexpression in dorsal CA1 introduced slow AMPA responses in a mosaic fashion. We therefore conclude that cell-type and region specific expression of CNIH3 is a driver of subcellular slow AMPA expression in the hippocampus.

## Results

In our previous study (Pampaloni et al., 2021), in addition to slow AMPA currents which developed and decayed over hundreds of milliseconds, we detected many cells in CA1 pyramidal cells that display only fast classical AMPA receptor synaptic currents with ∼10 ms decays. This cell-to-cell variability provides an obvious explanation for the general lack of reports of slow AMPA in the literature. Dentate granule cells also completely lack slow AMPA. Noting that not only are CA1 pyramidal cells inherently diverse, but that this diversity is expressed along the “natural” axes of the hippocampus (Cembrowski et al., 2016b), we hypothesised that slow AMPA could have regional expression in the mouse hippocampus. We divided organotypic slices between dorsal and ventral regions (Fig. 1A), and performing patch clamp on pyramidal cells in CA1 and granule cells in DG, we subjected dendritic regions to 10 Hz MNI glutamate uncaging (Fig. 1B). We blocked NMDA, kainate and GABA-A receptors and used near-physiological concentration of magnesium (1 mM). As we already reported (Pampaloni et al., 2021), the remaining currents activated by MNI-glutamate uncaging in CA1 pyramidal neurons at –60 mV rely on AMPAR conductances, because the application of GYKI 52466 (AMPAR blocker) abolished them (Fig. 1. Supp. 1.). However, as expected, we observed an incomplete reduction in CA3 pyramidal neurons, consistent with the involvement of kainate-type glutamate receptors resistant to UBP 310, and did not explore AMPAR kinetics in this subregion (Fig. 1. Supp.1). We defined slow AMPA responses as inward currents that displayed, in addition to a fast activating current, an accumulating slow response at 10 Hz of more than 10% of the original peak current, that had a weighted decay constant > 25 ms after the train of 10 pulses of glutamate ended (Fig. 1B). Using these dual criteria (decay time constant and *I*_ss_) to define slow AMPA lessened the chance of false positives caused by slow drifts in the current baseline. Moreover, we observed a substantial variability of the response amplitudes, ranging from 10 to almost 400 pA measured on the first peak of the 10 pulse train (Fig. 1 Supp. 2). Close inspection of the path of the UV pulse revealed that the highest amplitudes correspond to a combination of 2 or more dendrites being stimulated. One-photon uncaging is relatively imprecise in the z-axis (Ellis-Davies, 2018), and so, aiming to accurately resolve responses from single sites, we thresholded the selected responses to a maximum of 100 pA (Fig. 1 Supp. 2). We likewise performed strict quality control on our recordings and included here only cells with non-damaged and cell-typical dendritic structure revealed by Atto-594 dye filling and displaying a R_access_ <50 MΩ. Any recordings with >200 pA leak were discarded.

**Figure 1.**
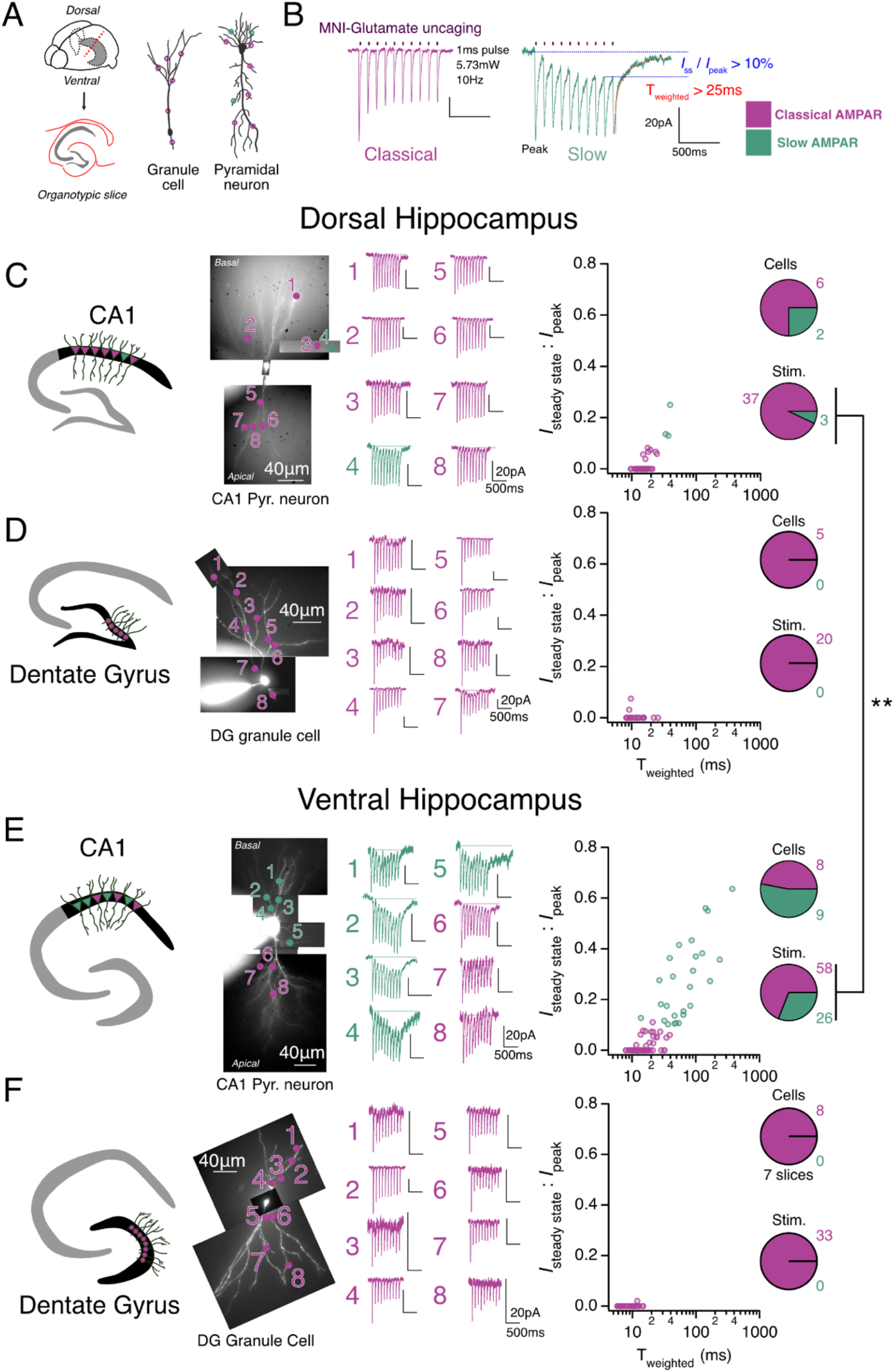
Slow AMPA is enriched in the ventral CA1 region of the mouse hippocampus. ***A:*** *Left:* Representative illustrations of hippocampus localization in the mouse brain, from which organotypic slices are dissected. *Right:* Cartoons of the neuronal architecture of granule cells of the Dentate Gyrus (DG) and Pyramidal neurons of CA1. 405 nm laser pulses (1 ms) were applied over the dendritic trees to uncage MNI-glutamate (sites indicated with circles). ***B:*** Protocol used for MNI-glutamate uncaging on recorded neurons in whole-cell patch clamp. 10 UV pulses at 10 Hz were applied to multiple dendrites in series in a single whole cell patch clamp recording. Dotted blue lines show the steady state current (*I*_ss_) and dotted red line the decay measured from the 10^th^ response (T_weighted_). Examples of typical classical (*I*_ss_ : *I*_peak_ < 0.1 and T_weighted_ < 25ms, purple) and slow (*I*_ss_ : *I*_peak_ > 0.1 and T_weighted_ > 25ms, green) responses are displayed. **C, D, E, F:** Measurement of AMPA kinetics in excitatory cells of Dorsal (C and D) and Ventral (E and F) mouse hippocampus. *Left:* Cartoon representation of the neuronal population recorded. *Middle*: Representative result of the responses recorded for 8 stimulation locations numbered on the tiled fluorescence micrograph, together with the associated currents, colour coded purple for classical fast AMPA and green for slow AMPA. *Right:* Two-dimensional plots with I_ss_ normalized to the 1^st^ peak response (*I*_ss_ : *I*_peak_) plotted with the associated T_weighted_ of the last (10^th^) response, where 1 point = 1 stimulation. Slow AMPA responses should lie in the upper right quadrant. Pie charts indicate the proportions of classical and slow cells (top) and responses (bottom), where a cell with at least one slow response is considered as slow. The incidence of Slow AMPA responses was strongly enriched in Ventral CA1 compared to dorsal CA1 (dCA1: 7.5% vs vCA1: 31%, *Pr of no difference* = 0.0032, Fisher’s exact test). dCA1: 8 cells / 5 slices, vCA1: 17 cells / 11 slices, dDG: 5 cells / 3 slices, vDG: 8 cells /7 slices.

We observed that under these conditions, slow AMPA responses are very rare in dorsal CA1 (3/40 sites, Fig. 1C). As previously reported, slow AMPA was completely absent in DG granule cells (Fig. 1D). However, strikingly, slow AMPA was prevalent in ventral CA1 pyramidal cells, with more than half the cells patched (9 out of 17) having slow responses (Fig. 1E). Overall, in ventral slices, about ⅓ of the responses from CA1 pyramids were slow (26/84 uncaging sites from 17 pyramidal cells). Within cells that showed a slow AMPA current, about half the responses were slow and half had a classical fast appearance. Despite a more stringent classification, the slow time constant of the decay for these slow responses was indistinguishable from our previous report, 182 ± 28 ms (40 ± 4% of the amplitude, *n* = 26 stimulation trains at 10 Hz), about 20-fold slower than a classical fast AMPA decay. Also, no difference of response amplitude was observed between the dorsal and ventral slices of the same subregion (namely CA1 or DG, Fig. 1 Supp. 3), and slow AMPA responses were still recorded in the same proportion when blocking K^+^ and Na_v_ channels with Cs-based intracellular solution and extracellular TTX respectively (Fig. 1 Supp. 4). Therefore, neither exaggerated response amplitudes, any eventual space clamp errors induced by K^+^ conductance, or a variation of slice activity with time or age (tested in our previous report, (Pampaloni et al., 2021), are likely to account for the kinetic variations observed in our recordings. These experiments demonstrate another reason why slow AMPA could be easily missed in the hippocampus, namely the easier access of the dorsal hippocampus could have led investigators to focus on this region. Related to this point, most studies of the hippocampus do not report whether they selected dorsal, ventral, central, or any combination of these subregions, in their work.

We previously demonstrated that slow AMPA responses could be evoked from Schaffer Collateral stimulation and miniature current durations were longer in slow AMPA positive neurons (Pampaloni et al., 2021). If the slow AMPA responses found from uncaging of glutamate come from receptors located at synapses, then electrical stimulation of the Schaffer collateral should also generate slow AMPA currents, and in proportion to the occurrence of slow AMPA in uncaging. We used the increased incidence of slow AMPA in ventral CA1 to test this idea. We mapped glutamate-triggered dendritic responses at 5 or more sites per pyramidal neuron, and stimulated in the stratum radiatum (Fig. 2). Perhaps unsurprisingly, cells in which only fast responses could be detected by uncaging gave classical fast EPSCs when the Schaffer Collateral was simulated, recapitulating hundreds of published results, and representing 9 out of 18 cells (Fig. 2A, 2B). The weighted time constant of decay after 1 s of 10 Hz stimulation of the SC was 18.8 ± 2.1ms. As the incidence of slow AMPA responses detected by uncaging increased, the response evoked by Schaffer stimulation had a slower decay and showed a larger steady-state current in response to repetitive stimulation (Fig. 2A). As previously reported (Pampaloni et al., 2021), in two cases where the recording could be held long enough, these slow SC responses were fully blocked by GYKI 52466, indicating that they arise from AMPA receptors (Fig. 2A). The Pearson *R* between the mean decay of responses to a train of 10 Hz uncaging and the mean decay from 10 Hz electrical stimulation in the same cell was 0.64 (Fig. 2C). The slope of the relation was 0.13, reflecting that Schaffer stimulation recruits a mixture of synapses, and the current response is the convolution of presynaptic adaptation and the post-synaptic short-term plasticity. However, these results suggest that the incidence of slow AMPA detected from uncaging reflects well the presence of slow AMPA synaptic currents in ventral CA1, readily activated by normal glutamate release with a modest 10 Hz stimulation.

**Figure 2.**
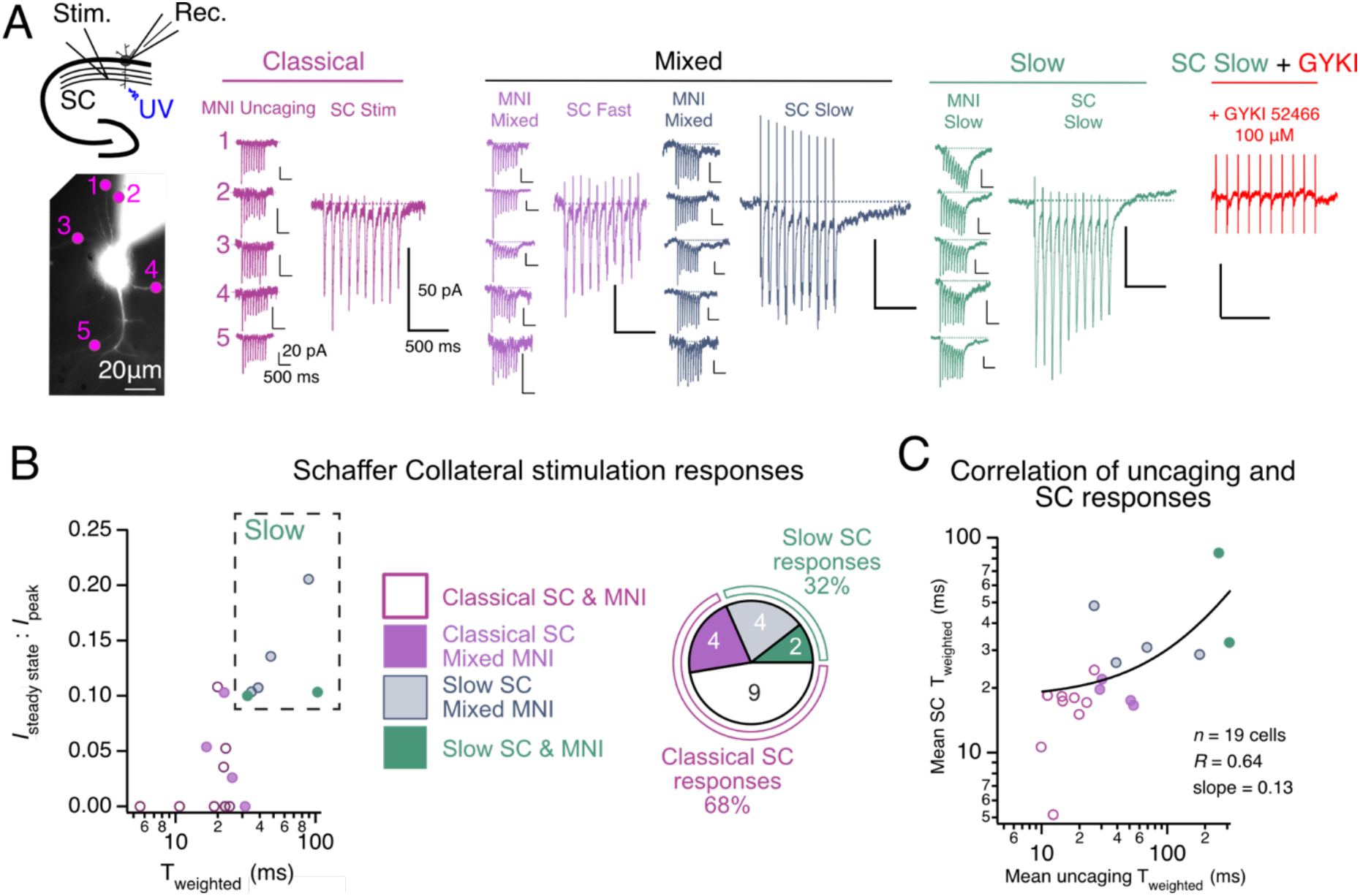
Electrical stimulation of identified slow AMPA-positive pyramidal cells. **A.** Recording of AMPA kinetics in ventral CA1 using electrical Schaffer Collateral (SC) stimulation and MNI-glutamate uncaging on the same cell. *Left:* illustration of the approach (Upper panel) and example of the fluorescence micrograph of a pyramidal neuron that showed only classical responses (bottom). *Middle:* Representative cells displaying different patterns of responses to MNI-glutamate uncaging and SC stimulation. Classical (classical MNI-uncaging and SC response); Mixed (mixed MNI-uncaging (*i.e* classical and slow) with Fast SC responses, or mixed MNI-uncaging with slow SC responses; and Slow (slow responses from MNI-glutamate uncaging and SC stimulation). *Right*: GYKI 52466 (100µM) inhibits a slow SC response completely (performed on *n* = 2 cells). **B.** Kinetics of SC stimulation responses. *Left*: Two-dimension plot with I_ss_ normalized to the 1^st^ peak response (*I*_ss_ : *I_peak_*) plotted with the associated T_weighted_ of the last (10^th^) response. Multiple protocols of SC stimulations were recorded on the same cell, and the slowest responses were selected (1 point = slowest SC decay for each cell). The color code indicating from which class of cell the response was recorded is shown on the side. The dotted rectangle highlights the SC slow responses. *Right*: Pie chart showing the proportion of the different patterns of responses. 68% of responses were classical SC, and 32% were slow SC. **C.** Correlation of MNI-uncaging and SC responses. The mean T_weighted_ of the SC responses (average of all the SC protocol stimulations performed on the same cell), and the mean T_weighted_ responses of the MNI-uncaging responses (average of all the dendrites stimulated on the same cell) were plotted together. 1 point = 1 cell. The colors are the same as B. For B and C, recordings were performed on *n* = 19 cells from 10 ventral slices.

Armed with the knowledge that slow AMPA synaptic currents are enriched in ventral CA1 and absent in dorsal CA1 and DG, we hypothesized that region-specific expression of certain AMPA receptor components could be responsible for the functional gradient of slow AMPA. We envisaged either enrichment of “positive” modulatory elements, which slow AMPA deactivation and reduce desensitization, or, in areas that lack slow AMPA, an enrichment for presumptive negative modulators. We examined published RNAseq databases for possible candidate molecules that could generate a slow AMPA receptor. We used the freely available dataset from the Janelia Research Campus reporting single-cell RNA expression of AMPAR complex transcripts in excitatory neurons of adult mice hippocampus (Cembrowski et al., 2016a) (Fig. 3). Other published datasets generated by other means (Allen Institute, Fig. 3. Supp. 1, (Yao et al., 2021) largely corroborate this analysis. We focused our analysis on the components of the native AMPAR complex in the rodent hippocampus (29 proteins revealed by mass spectrometry of GluA1 or GluA2 precipitated complexes, (Schwenk et al., 2014). The expression profiles were plotted to help identify preferential AMPAR components along the ‘long’ axis of CA1, whilst also acknowledging their absolute expression in CA1 and DG. Most AMPAR components determined from proteomics (20/29) show no appreciable expression gradient (< 3-fold difference) on the cellular level, but several presumptive positive and negative modulators of AMPA receptor activation do show appropriate differences. For example, CNIH3, GluA4 and γ-8 show strong enrichment gradients. However, γ-8 is known to be strongly expressed in DG, and overexpression of dominant negative γ-8 did not change slow AMPA incidence much (Pampaloni et al., 2021). In CA1, KO of γ-8 reduces fast AMPA synaptic currents by 35%, and extrasynaptic currents by ∼90% (Rouach et al., 2005) consistent with major roles in trafficking and anchoring receptors at synapses (Watson et al., 2021; Sheng et al., 2018). Based on single channel recordings (Zhang et al., 2014), we consider GluA4 as a putative slow subunit, even though it is largely considered to be a very fast AMPAR subunit expressed in interneurons (Geiger et al., 1995). GluA4 shows appreciable transcript enrichment in ventral CA1 pyramidal cells, but it remains unclear whether it is actually expressed there (based on the lack of synaptic currents in the GluA123 KO mouse (Lu et al., 2009)). On the other hand, negative modulators of AMPA gating including the auxiliary subunits Shisa9 (also called CKAMP44) and GSG1-L are enriched in the DG, potentially contributing to the suppression of slow AMPA there (von Engelhardt et al., 2010; Gu et al., 2016). Even though CNIH3 is shown to play a role in hippocampus-related spatial memory (Frye et al., 2021; Lintz et al., 2026), its effect on AMPAR kinetics beyond heterologous expression (Schwenk et al., 2009) (Shanks et al., 2014) remains unclear (Bharti et al., 2025; Herring et al., 2013). Bearing in mind contradictory reports about surface expression of Cornichons and its modulation of kinetics (Coombs et al., 2012) we reprised measurements in HEK 293 cells to see whether CNIH3 produced receptors with the correct activation profile for slow AMPA.

**Figure 3.**
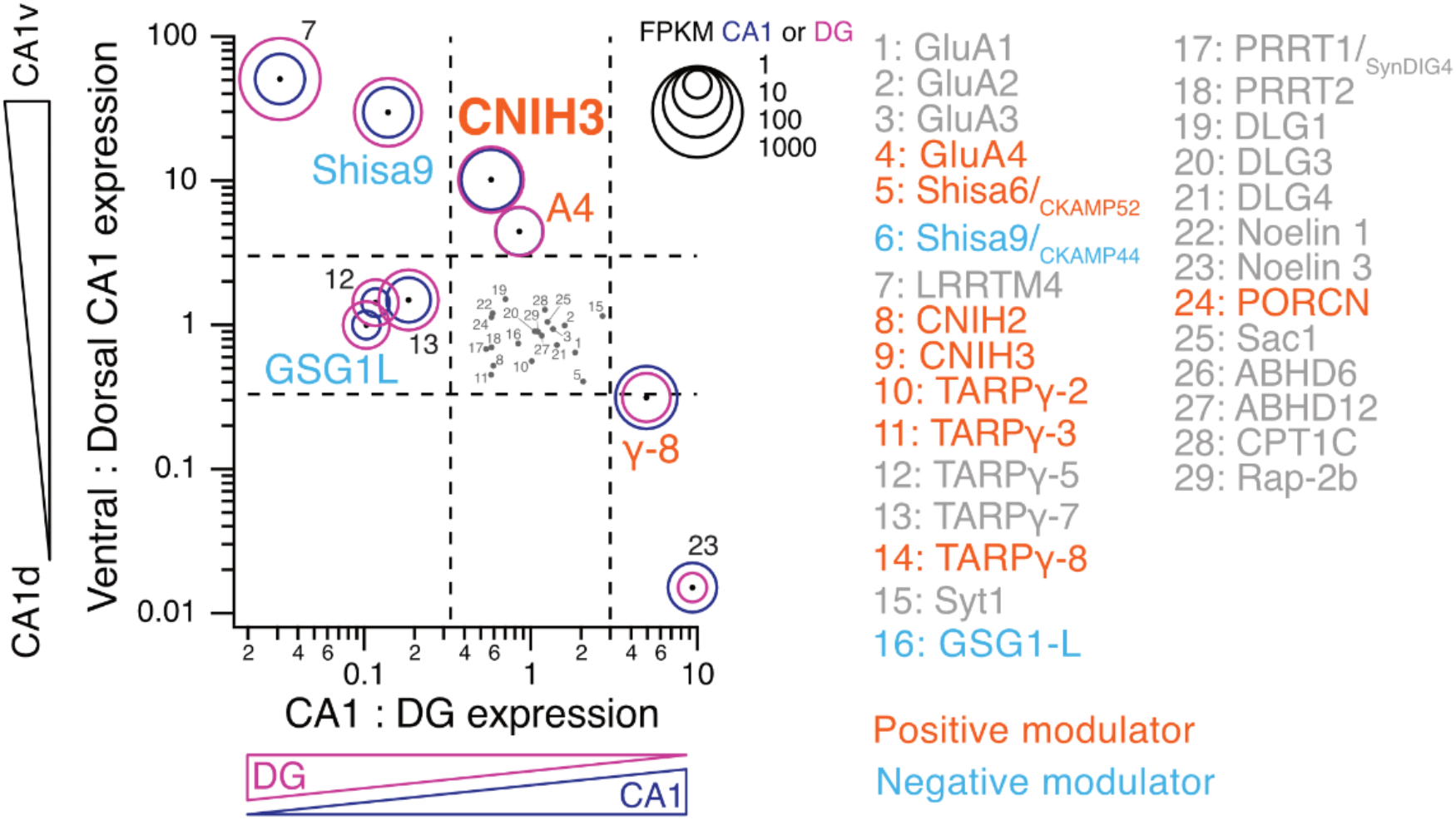
Identifying candidates for slow AMPA from gradients of AMPAR components in single cell transcriptomics from the mouse hippocampus. Single-cell RNA expression of AMPAR complex transcripts in excitatory neurons of adult mice hippocampus. Fragments Per Kilobase Million (FPKM) values from the HippoSeq data set from ventral and dorsal hippocampus using microdissected CA1 and Dentate Gyrus (DG). https://hipposeq.janelia.org,(Cembrowski et al., 2016a). The selected transcripts compose native AMPAR complexes from mass spectrometry of rat brain (Schwenk et al., 2014), and the numbers in the graph correspond to the list on the side. The diameters of the circles indicate the abundance of each transcript in CA1 (blue) and DG (purple) (see the inset scale for FPKM), and the points at their centres are the ratios of the FPKMs of Ventral and Dorsal CA1 (Y axis) is plotted together with the ratio of the FPKMs of CA1 and DG (X axis). The dotted lines indicate the threshold for a meaningful difference (more than 3-fold enriched or depleted). The presumptive positive (orange) and negative (blue) regulators of AMPA activation are indicated (orange: promoting activation, blue: reducing activation). CNIH3 is highlighted as the main candidate (known to slow down AMPA decay and enriched in ventral CA1).

Fast perfusion of glutamate onto outside-out patches pulled from HEK 293 cells expressing homomeric and heteromeric AMPA receptors in the presence and absence of CNIH3 revealed a near universal and profound slowing of AMPAR kinetics, and resistance to desensitisation (Figure 4), similar to previous reports (Schwenk et al., 2009; Shanks et al., 2014; Miguez-Cabello et al., 2025). CNIH3 expression was also highly deleterious for cell health. Some previous studies focused on desensitization kinetics (Hawken et al., 2017), which is less important for synaptic receptors. We instead concentrated on resistance to desensitization and slow deactivation kinetics (either after long or short glutamate applications) which are relevant for synaptic currents, and are needed for slow AMPA responses.

**Figure 4.**
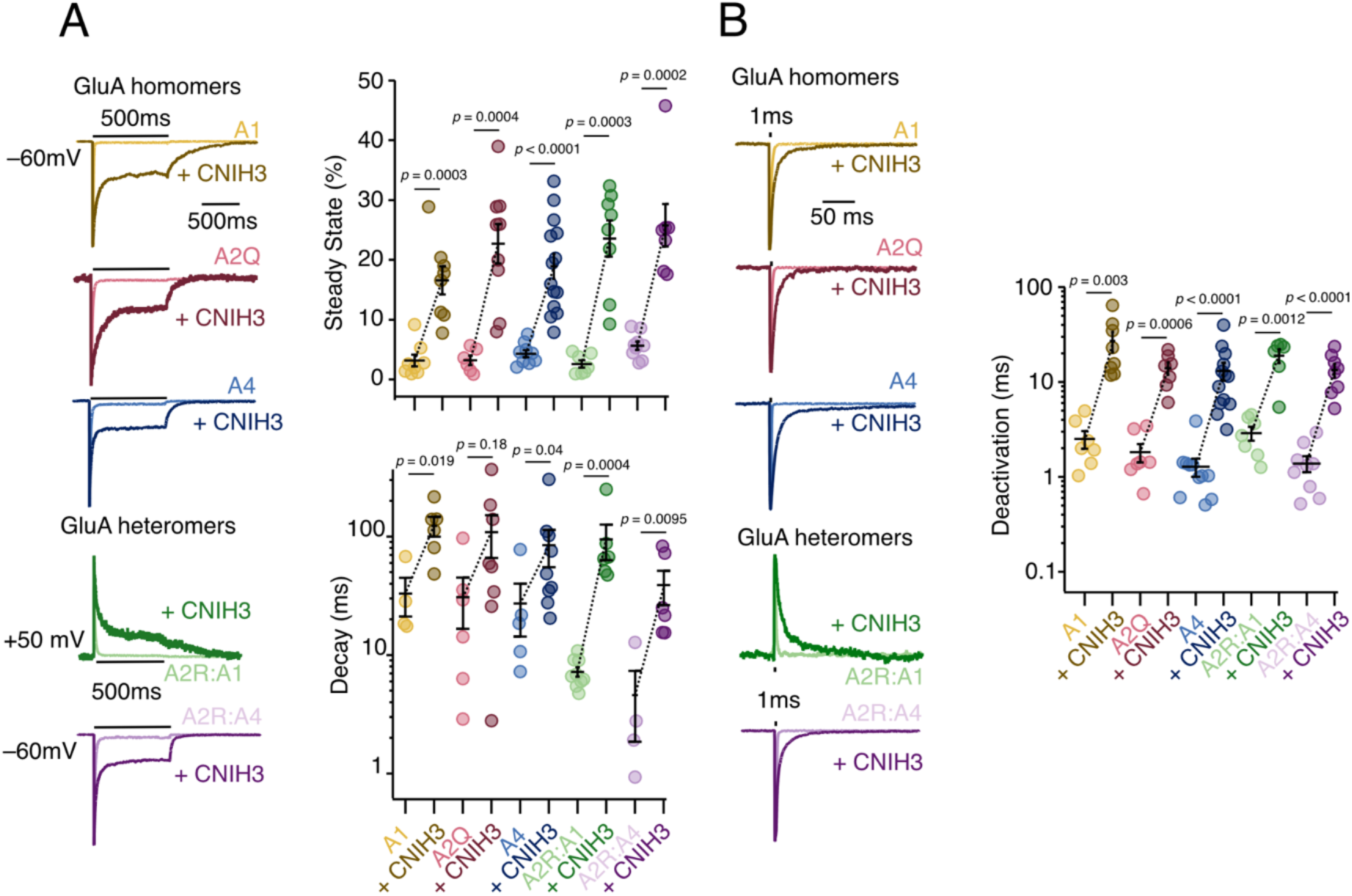
CNIH3 is a strong modulator of AMPA kinetics. **A.** Effect of CNIH3 on AMPAR kinetics during a long (500 ms) glutamate exposure. *Top Left*: representative recordings of AMPA homomeric receptors with (dark traces) and without (light traces) CNIH3 co-expression. *Bottom Left*: representative recordings of GluA heteromer receptors with (dark traces) and without (light traces) CNIH3 co-expression. *Top Righ*t: Steady state (%) is significantly increased for all homomeric and heteromeric AMPARs in the presence of CNIH3. Dashed lines indicate the changes in the mean. For each condition respectively: GluA1 (from 3.2 ± 1.0% to 16.6 ± 2.4%, p = 0.0003), GluA2Q (from 3.2 ± 0.8% to 22.7 ± 3.3%, p = 0.0004), GluA4 (from 4.3 ± 0.6% to 18.9 ± 2.1%, p < 0.0001), GluA2R:1 (from 2.6 ± 0.6% to 23.6 ± 3.0%, p = 0.0003), and GluA2R:4 (from 5.7 ± 0.7% to 25.8 ± 3.6%, p = 0.0002), without and with CNIH3.. *Bottom Right*: Receptor decay is substantially slower for GluA1 and GluA4 homomers and GluA1:GluA2R and GluA4:GluA2R heteromers when co-expressed with CNIH3; GluA1 (from 33.1 ± 11.9% to 123.4 ± 23.7%, p = 0.019), GluA2Q (from 30.9 ± 14.2% to 109 ± 43%, p = 0.18), GluA4 (from 27.3 ± 13% to 84.6 ± 29.4%, p < 0.04), GluA2R:1 (from 7.2 ± 0.7% to 94.9 ± 31.7%, p = 0.0004), and GluA2R:4 (from 4.6 ± 2.8% to 39 ± 12.5%, p = 0.0095), without and with CNIH3, respectively. Error bars represent standard error of the mean. **B.** Effect of CNIH3 on different AMPA receptor deactivation during a short (1 ms) glutamate exposure. *Top Left*: representative recordings of GluA homomeric receptors with (dark traces) and without (light traces) CNIH3 co-expression. *Bottom Left:* representative recordings of AMPAR heteromers with (dark traces) and without (light traces) CNIH3 co-expression. *Right*: Deactivation of different AMPA receptors is universally slower in the presence of CNIH3, GluA1 (from 2.5 ± 0.5% to 27.2 ± 6.6%, p = 0.003), GluA2Q (from 1.8 ± 0.4% to 14.02 ± 2.1%, p = 0.0006), GluA4 (from 1.3 ± 0.3% to 13.1 ± 2.8%, p < 0.0001), GluA2R:1 (from 2.9 ± 0.5% to 19 ± 3.1%, p = 0.0012), and GluA2R:4 (from 1.4 ± 0.3% to 13.4 ± 2.2%, p < 0.0001), without and with CNIH3, respectively. Error bars represent standard error of the mean. Statistical significance was tested using the Mann-Whitney test.

Without CNIH3, the responses in outside-out patch recapitulated the classical fast AMPA responses. In the presence of CNIH3, AMPA receptors still showed fast activation, but we observed strong block of desensitisation in GluA1, GluA2(Q), and GluA4 homomers, as well as GluA2R:A1 and GluA2R:A4 heteromers (Fig. 4 Supp. 1). The steady-state current was in the range 15-25% in response to a 500 ms pulse of 10 mM glutamate (Fig. 4A). The decay time constant after the glutamate was removed, akin to the current decay after a burst of stimulation of synaptic receptors, was in the 100 ms range, similar to (but less than) what we observed in ventral CA1 (see Fig. 1E). For heteromeric receptors, the slowing of the decay due to CNIH3 was more profound, about 20-fold, than in homomeric receptors (average T_weighted_ decay for homomeric channels without CNIH3 was 27 - 33 ms and increased to 84 −123 ms with CNIH3 co-expression, while average T_weighted_ decay for heteromeric complexes in the absence of CNIH3 was 4-7 ms, and increased to 38 - 94 ms with CNIH3). Likewise, in response to a 1 ms pulse of glutamate, the deactivation were universally slowed by CNIH3, with average T_weighted_ time constants ranging from 1.3 - 2.5 ms for AMPA receptors alone and 13 - 27 ms for receptors co-expressed with CNIH3 (Fig. 4B). Given recent structural results that place CNIH proteins firmly within hippocampal AMPA receptor complexes (Yu et al., 2021), we expect that the receptors expressed on the HEK cells surface are complexes between GluA subunits and CNIH3, which have quite similar kinetics to the responses seen in ventral CA1. To more closely mimic the hippocampal slice uncaging experiments, we measured currents from outside-out patches of HEK cells expressing a heteromeric GluA2R:A4 receptor, with or without CNIH3 co-expression, to 1 ms train pulses at frequencies ranging from 10 - 50 Hz (Fig. 4 Supp. Fig. 2). Without CNIH3, there was no steady state current. At 10 Hz, inclusion of CNIH3 slowed the receptor decay, partially reproducing the kinetics observed in neurons (almost reaching the 25 ms / 10% I_ss_/I_peak_ threshold). As expected, with higher frequencies, a more profound slowing of receptor kinetics was measured in the presence of CNIH3. A synapse with CNIH3-containing receptors with high frequency activity, could carry exaggerated charge transfer from the steady state current together with slow decay at the end of the train. While the obtained kinetics were not as slow as those measured in slice recordings, these findings suggest that CNIH3 can endow receptors with kinetics similar to endogenous slow AMPARs.

The enrichment of CNIH3 in ventral CA1 and its capacity to generate AMPA receptors with slow decays and resistance to desensitization led us to hypothesise that it was a decisive component of slow AMPA expression in the ventral CA1. Knockdown of CNIH3 should therefore reduce the incidence of slow AMPA in ventral CA1. We obtained two separate shRNAs specific for CNIH3 (see Methods), which we expressed in mouse ventral organotypic slices by AAV transduction. A separate, scrambled shRNA was used as a control. We measured the incidence of slow AMPA using 10 Hz glutamate uncaging in patch clamp as before (Fig. 5). Experiments were carried out blind to the shRNA and transduced cells were identified from their GFP expression (Fig. 5A). Strikingly, both shRNAs against CNIH3 abolished slow AMPA responses for all practical purposes (1 slow AMPA response from 179 stimulations across the two shRNAs, mean decay of 13.8 ± 0.4 ms for shRNA_1 and 14.4 ± 0.8 ms for shRNA_2, Fig. 5C, 5D). In contrast, slow AMPA was retained in slices that were transduced with the scrambled shRNA, albeit with a mildly reduced incidence (mean decay of 18.5 ± 1.5 from 145 stims, including 12 slow AMPA responses, Fig. 5B). We observed no difference in the amplitude of the responses, suggesting the KD of CNIH3 does not affect the AMPAR expression or dendritic targeting (Fig. 5 Supp. 1). This result strongly suggests that CNIH3 expression is needed for slow AMPA.

**Figure 5.**
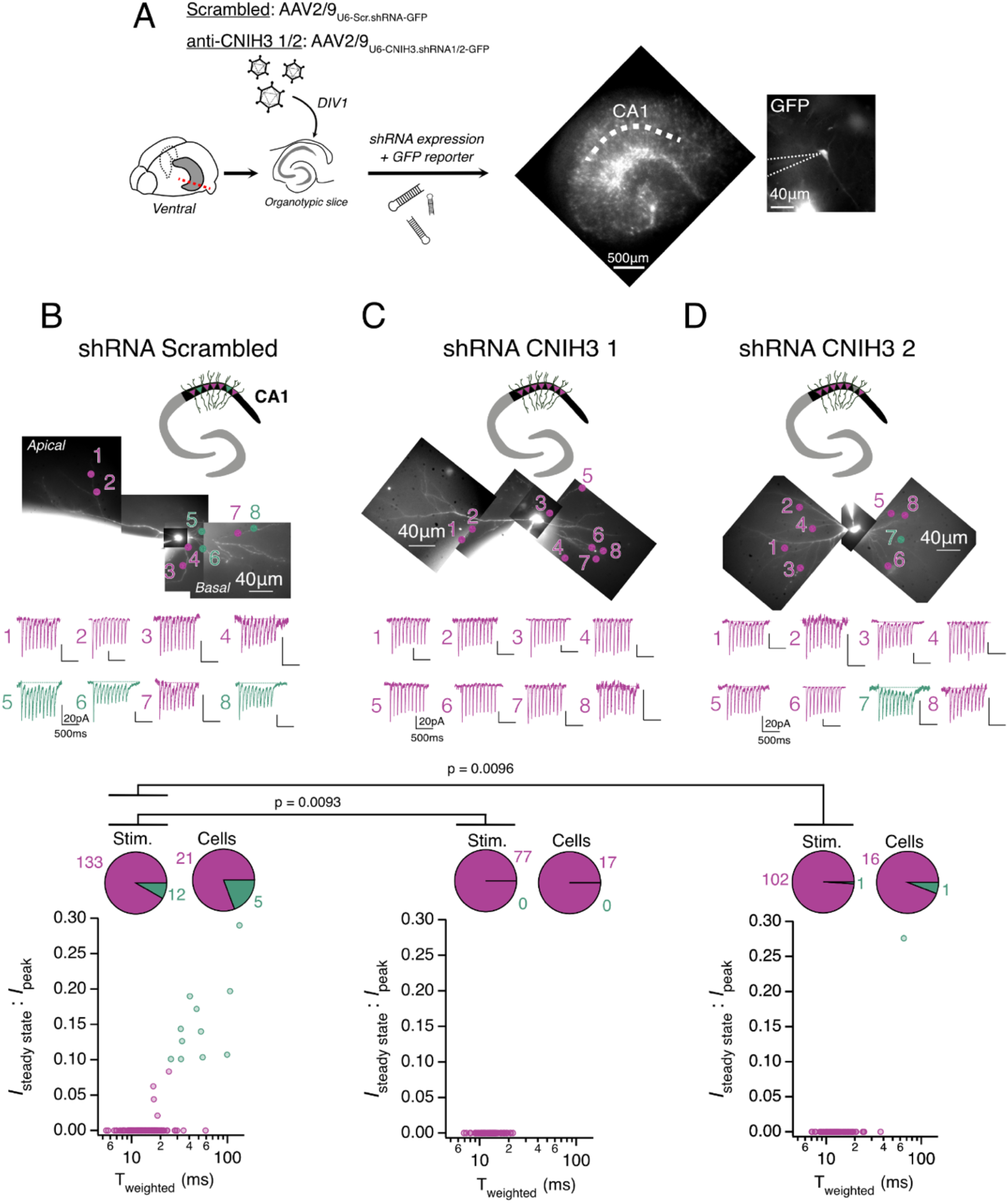
Knockdown of CNIH3 abolishes slow AMPA in ventral CA1. ***A:*** *Left:* Graphical illustration of the CNIH3 knock-down protocol. AAV-Scrambled or AAV-ShRNA CNIH3 1 or 2 were applied at DIV 1 on mouse hippocampus organotypic slices. *Right:* AAV-transduced cells express GFP, as depicted in the representative fluorescence of a ventral organotypic slice (4 x objective); and of a pyramidal neuron on the side (63 x objective) where the recording pipette is highlighted by dotted lines. ***B, C and D:*** Measurement of AMPA kinetics in response to glutamate uncaging in pyramidal neurons expressing shRNA scrambled (B), shRNA CNIH3 1 (C) and shRNA CNIH3 2 (D) in ventral CA1. *Up:* Cartoon representation of the neuronal population recorded (purple: slow, and green: classical), together with a representative result of the responses recorded with 8 stimulations, with their positions marked on the tiled fluorescence micrograph. *Bottom:* Two-dimension plots with *I*_steady state_ normalized to the 1^st^ peak response (I_ss_ / I_peak_) plotted with the associated T_weighted_ of the last (10^th^) response, where 1 point = 1 stimulation. Pie charts indicate the proportion of classical and slow cells & stimulations, where a cell with at least one slow response is considered as slow. The incidence of Slow AMPA responses was strongly reduced by CNIH3 shRNA compared to the Scr. shRNA (Scr: 8%) vs anti-CNIH3_1 (0%, *Pr of no difference* = 0.0093, Fisher’s exact test) and vs anti-CNIH3_2 (1%, *Pr of no difference* = 0.0096, Fisher’s exact test). Bonferroni corrected ɑ = 0.025 (0.05/2). Scr: 26 cells / 14 slices, shRNA_1: 17 cells / 10 slices, shRNA_2: 17 cells / 8 slices.

Emboldened by this unusually clear result, we next asked if the absence of CNIH3 transcripts in dorsal CA1 was a factor stopping slow AMPA expression there. We compared the incidence of slow AMPA in slices where we overexpressed CNIH3 from an AAV, to control slices where we expressed GFP (also from AAV, Fig. 6A). These experiments were not done blind because the GFP expression level was so different between the two conditions, and the overexpression of CNIH3 was generally not well tolerated by the slices (see discussion). Nonetheless, the incidence of slow AMPA was considerably higher in CNIH3-expressing neurons than in control conditions (24/122 in CNIH3 overexpression vs 8/110 in GFP controls, *p* = 0.0073, Fig. 6B, 6C). The level of slow AMPA in GFP controls was indistinguishable from that in our original recordings of non-transfected wild-type slices (see Fig. 1C). In contrast to previous experiments with overexpression of γ-8 in CA1 (Pampaloni et al., 2021), whereby all uncaging sites developed a large slow AMPA current, completely different from the physiological ⅓ incidence, for CNIH3 overexpression, the physiological mosaic expression of slow AMPA was recapitulated with 6 out of 18 transduced cells still only showing classical fast responses. The remaining 12 cells showed slow AMPA (74 ± 14ms decay and *I*_ss_ : *I*_peak_ = 28 ± 5 %) at some uncaging sites and also showed typical fast classical AMPA responses at other sites (Fig. 6C). When we overexpressed CNIH3, the proportion of slow responses in dorsal CA1 were not significantly different from the proportion we found in WT ventral CA1 (p = 0.07, Fisher’s exact test, see Fig. 1E). These results suggest that CNIH3 expression is enough to introduce slow AMPA but that other factors (either synaptic or non-synaptic) decide where the slow AMPA current is expressed or not.

**Figure 6.**
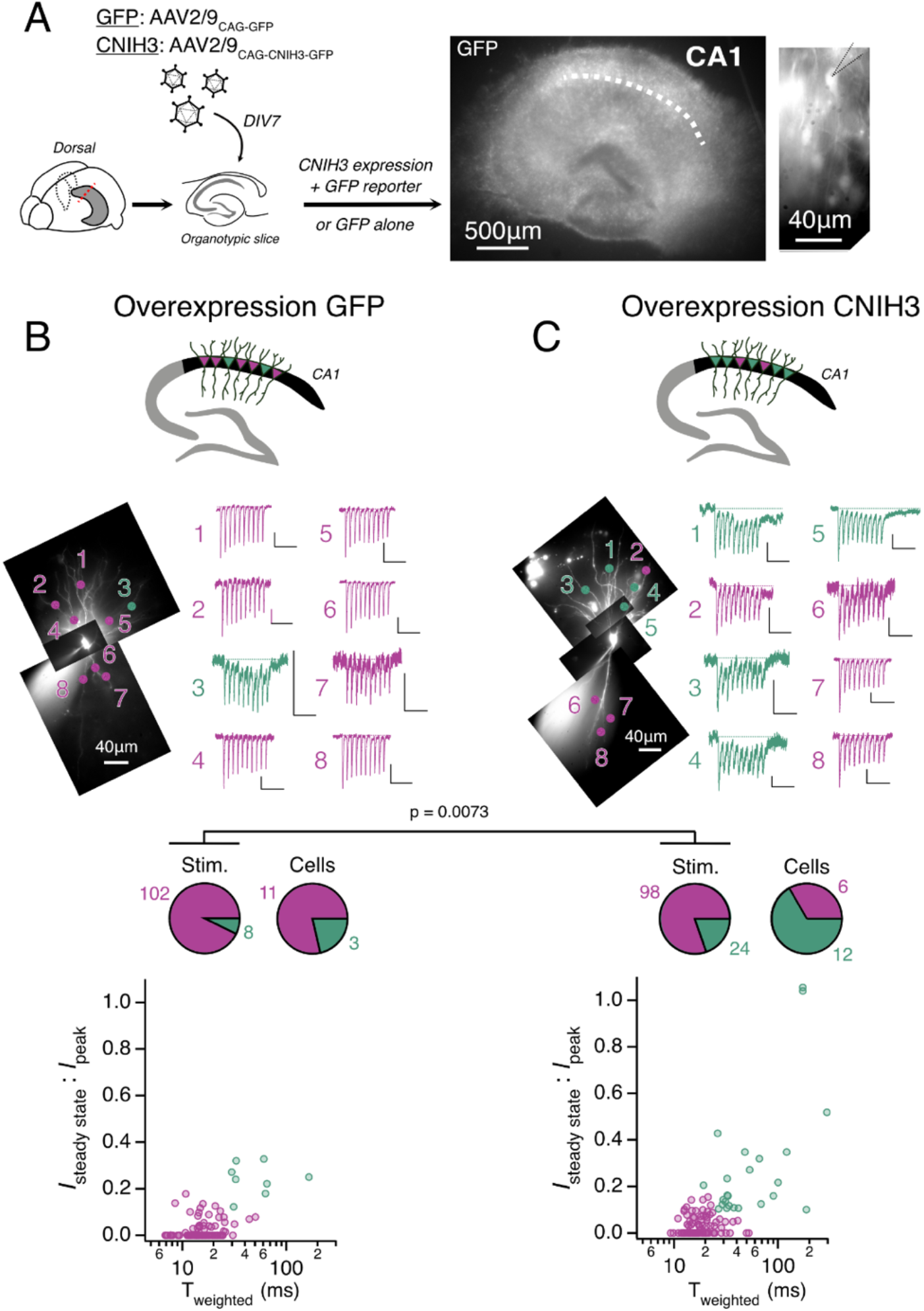
Overexpression of CNIH3 introduces mosaic slow AMPA responses in dorsal CA1. ***A:*** *Left:* Graphical illustration of the CNIH3 or GFP overexpression protocol. AAV-GFP or AAV-CNIH3 were introduced to mouse hippocampus organotypic slices on DIV 7. *Right:* AAV-transduced cells express GFP, as depicted in the representative fluorescence of a ventral organotypic slice (4x objective); and of a pyramidal neuron on the side (63 x objective) where the recording pipette is indicated by dotted lines. ***B and C:*** Measurement of AMPA kinetics in response to glutamate uncaging in pyramidal neurons expressing only GFP (B) or CNIH3 and GFP (C) in dorsal CA1. *Upper:* Graphical representation of the neuronal population recorded (purple: slow, and green: classical), together with a representative result of the responses recorded with each time 8 stimulations localized on the tiled fluorescence micrograph. *Lower:* Two-dimension plots with *I*_steady state_ normalized to the 1^st^ peak response (I_ss_ / I_peak_) plotted with the associated T_weighted_ of the last (10^th^) response, where 1 point = 1 stimulation. Pie charts indicate the proportion of classical and slow cells & stimulations, where a cell with at least one slow response is considered as slow. The incidence of slow AMPA responses at individual sites was strongly increased by the CNIH3 expression compared to the GFP expression alone. GFP (7.2 %) vs CNIH3 1 (19,6%, *Pr of no difference* = 0.0073, Fisher’s exact test). GFP: 14 cells / 7 slices, CNIH3: 18 cells / 11 slices.

Given that auxiliary subunits change glutamate receptor pharmacology, and we have relied on GYKI 52466 to confirm the identity of AMPA receptors in the hippocampus (see (Pampaloni et al., 2021), we used heterologous expression in HEK 293 cells to confirm that CNIH3-containing receptors retain sensitivity to GYKI 52466. In outside out patches, GYKI 52466 inhibited glutamate-activated peak currents (Fig. 7A-B), but was less effective on steady state currents, and in particular the steady state response of GluA4-CNIH3 seemed somewhat resistant to GYKI (from 17 ± 2.8% GluA4 alone vs 28 ± 1.8% with CNIH3, despite a reduction in the absolute current) (Fig. 7C).

**Figure 7.**
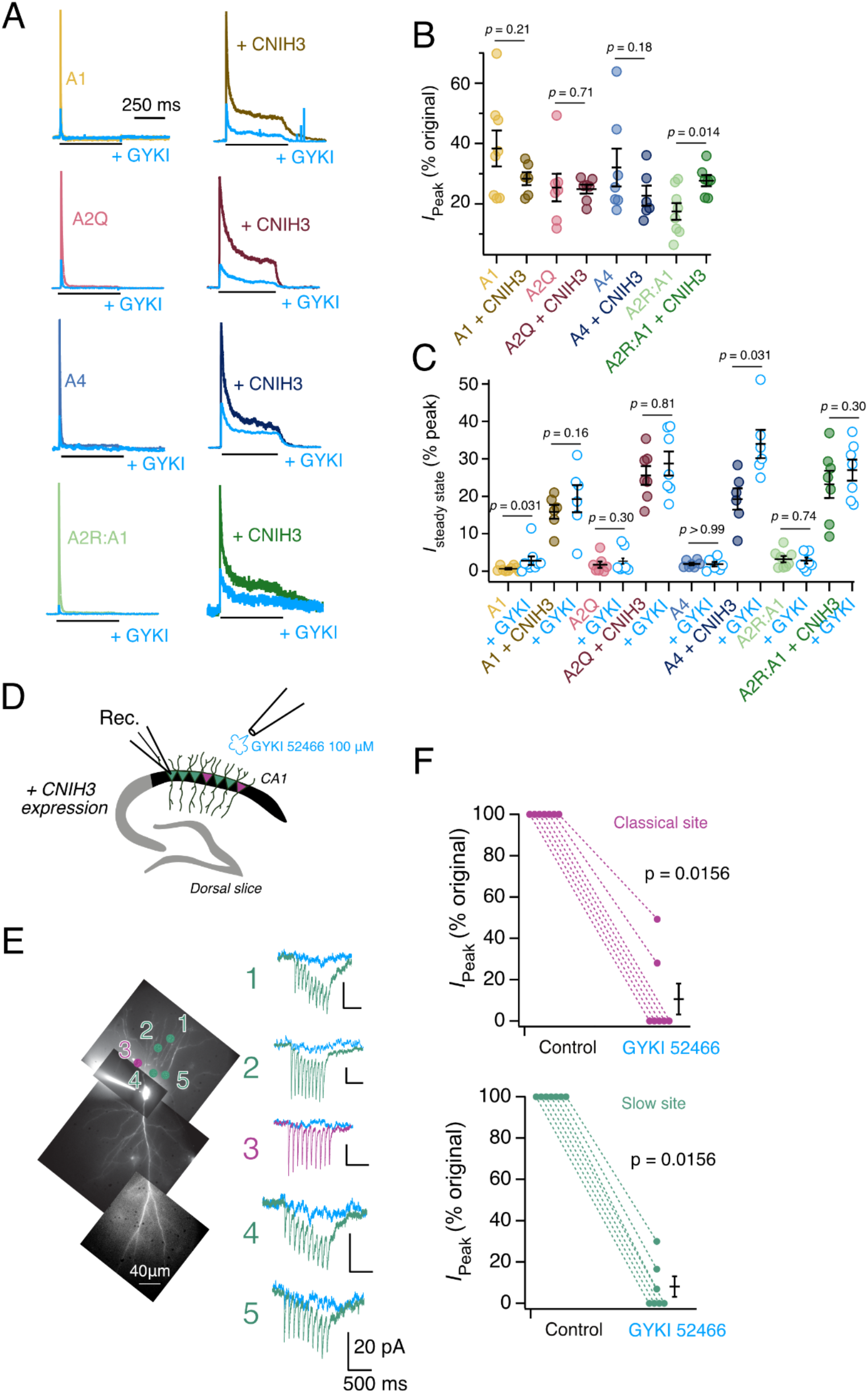
CNIH3 does not change AMPAR inhibition by GYKI 52466. **A:** Effect of CNIH3 on GYKI 52466 AMPA receptor response. *Left*: Representative recordings of GYKI inhibition of AMPA receptors in the absence of CNIH3 (light traces). *Right*: Representative recordings of GYKI inhibition of AMPA receptor current when co-expressed with CNIH3 (dark traces). In all conditions, application of GYKI 52466 (light blue traces) inhibits both peak and steady state of the current response. Recordings measured at +50 mV during a 500 ms application of either 10 mM glutamate or 10 mM glutamate + 100 μM GYKI 52466. **B:** Peak inhibition of GYKI 52466 on AMPA receptors with and without CNIH3. With the exception of GluA2R:1, the addition of CNIH3 does not change the peak inhibition of GYKI 52466; GluA1 (from 38.4 ± 5.9% to 28.4 ± 2.2%, p = 0.21), GluA2Q (from 25.4 ± 4.6% to 24.9 ± 1.4%, p = 0.71), and GluA4 (from 32.1 ± 6.3% to 22.7 ± 3.3%, p = 0.18), without and with CNIH3 co-expression, respectively. For GluA2R:1, the co-expression of CNIH3 decreased the inhibitory effect of GYKI 52466, with peak inhibition increasing from 17 ± 2.8% to 28 ± 1.8%. **C:** Steady state inhibition of GYKI 52466 on AMPA receptors, with and without CNIH3. In nearly all AMPA receptors, the addition of CNIH3 does not alter the steady state with GYKI 52466; GluA1 + CNIH3 (from 15.9 ± 1.9% to 19.3 ± 3.7%, p = 0.16), GluA2Q (from 1.8 ± 0.8% to 2.7 ± 1.2%, p = 0.30), GluA2Q + CNIH3 (from 25.6 ± 2.5% to 28.8 ± 3.2%, p = 0.81), GluA4 (from 1.9 ± 0.3% to 1.9 ± 0.6%, p > 0.99), GluA2R:1 (from 3.3 ± 0.8% to 2.9 ± 0.8%, p = 0.74), GluA2R:1 + CNIH3 (from 23.2 ± 3.6% to 27.0 ± 2.8%, p = 0.30), with glutamate and with glutamate + GYKI 52466, respectively. Note that the Steady State of GluA4 with CNIH3 is elevated during GYKI application (from 19 ± 2.9% to 34 ± 4 %) as well for GluA1 alone (from 0.7 ± 0.2% to 2.8 ± 1.1%, p = 0.031). Error bars represent standard error of the mean. **D:** Graphical illustration of the protocol, where MNI-glutamate uncaging on a CNIH3 overexpressing pyramidal neurons was performed in dorsal CA1. The same site was stimulated with or without GYKI 52466 (100 µM), an AMPAR-specific blocker, applied with a micropipette on the slice. **E:** Representative responses recorded from a dCA1 pyramidal neuron expressing CNIH3 (fluorescence tiled micrograph with stimulation localizations depicted) with or without GYKI 52466 (light blue). Purple: Classical response; Green: Slow response. **F:** GYKI strongly inhibits the classical *(Upper, 89* ± 7.5% reduction) and slow *(Lower, 92* ± 4.8% reduction) responses. Responses are plotted in % of the original *I*_peak_ response recorded before GYKI application. All the recordings were done in 4 cells on *n* = 14 sites (7 classical and 7 slow). All the statistics were done using the Wilcoxon matched-pairs signed rank test.

In dorsal CA1 pyramidal cells in which we overexpressed CNIH3, we checked to see that the slow responses from glutamate uncaging still showed the expected inhibition from GYKI 52466 (that is, that these responses were from AMPA receptors), Fig. 7D, E, F. There was no difference in the GYKI inhibition of slow or fast AMPA responses in these experiments (Fig. 7E, 7F), but the inhibition was generally stronger than in the HEK-293 experiments (73 ± 1.6% of inhibition in HEK-293 experiments, all subunits included vs 90 ± 4.3% of inhibition in neurons, all sites included). Given that TARPs γ-2 and γ-8 increase the sensitivity to GYKI (Pampaloni et al., 2021) these results, whilst confirming that CNIH3 expression drives slow AMPA to be expressed, suggest that other auxiliary proteins could be involved, in order that the very complete GYKI inhibition observed in CA1 pyramidal cell slow AMPA can be reconciled with the partly incomplete inhibition of receptors with and without CNIH3 in HEK cells.

## Discussion

In this work, we have identified a new actor in AMPA receptor and synapse diversity in the hippocampus. Knockdown and overexpression experiments based on transcriptomics and the functional gradient of slow AMPA responses indicate that CNIH3 is a decisive factor in producing slow AMPA currents. Our results clearly indicate an AMPA-receptor based mechanism for slow excitatory responses, distinct from other “slow” glutamate receptor types, be they ionotropic or metabotropic. Other elements are certainly required to support a stable distribution of slow AMPA responses across cell types and the mosaic subcellular distribution of positive and negative synapses along the same dendrites. But the experiments presented in this paper strongly suggest that CNIH3 is a component of non-desensitizing, slow AMPA receptors which produce substantial synaptic decays in the hundreds of milliseconds timescale, highly divergent from the canonical view of AMPA receptors as the fastest receptor in the brain.

Slow AMPA responses are concentrated in ventral CA1 (31% of the uncaging sites), rarely observed in dorsal CA1 (7.5% of the sites) and are absent in the dentate gyrus granule cells in both dorsal and ventral regions (Fig. 1). The bi-directional change in slow AMPA responses to glutamate driven by CNIH3, including potent knockdown in ventral CA1 by shRNA and introduction of slow responses to dorsal CA1 associated with a mosaic synaptic distribution, in overexpression, strongly suggests several facts. Slow AMPA responses depend on the molecular basis of AMPA receptors. Slow AMPA currents have a synapse-specific distribution, determined by synapse identity. Slow AMPA receptors are expressed in a cell specific fashion, even in ‘common’ cell types such as CA1 pyramidal neurons.

The role of the CA1 in rodents has been extensively explored over more than 50 years (O’Keefe, 1976), and its connectivity within the ventral region and with other brain areas is diverse (Montero-Crespo et al., 2020; Hong and Kaang, 2022). Accumulating work reveals a clear differentiation between ventral and dorsal poles of the CA1 in a variety of functions, such as goal-directed behavior (Hasantash et al., 2025) and contextual representation (Mishchanchuk et al., 2024). Accordingly, functional connectivity along the ‘long’ axis of CA1 has been reported (Hong and Kaang, 2022), together with gradients of electrophysiological properties (Malik et al., 2016) and gene expression (Cembrowski et al., 2016b) in pyramidal neurons. If we make the assumption that the slow AMPA synapses are confined in a neuronal subgroup, and not a spatially-undefined, diffuse property, we can hypothesise that this neuronal ensemble would have a role in a vCA1-related function(s). The involvement of vCA1 in contextual fear memory has been largely reported for instance (Hong and Kaang, 2022), involving specific ensembles of neurons encoding aversive stimuli (Jimenez et al., 2020; Gergues et al., 2020). The molecular mechanism underlying vCA1-based memory is still unclear.

As a mechanism of short term potentiation, slow AMPARs comfortably exceed the power of other post-synaptic mechanisms reported to date, and offer flexibility over presynaptic mechanisms like boosted release probability (which do not extend responses in time). Likewise, the mosaic expression of slow AMPA in single cells is more pronounced than previously reported receptor gradients or heterogeneity in dendrites (Andrasfalvy and Magee, 2001; Tóth and McBain, 1998). Given the mosaic distribution of slow AMPA, it is tempting to speculate that synapses might incorporate slow AMPA as a form of information storage, or as part of the mechanism of storage. Classical Hebbian spike-timing dependent plasticity works on the same timescale as fast AMPA receptors, but other forms of plasticity need slower time constants. Behavioural time scale plasticity (BTSP, (Bittner et al., 2017; Magee, 2026), which operates over about half a second, can drive one-shot learning through the CA1 network, but functions in dorsal hippocampus, which seems to lack slow AMPA. Even more intriguing are the detonator properties of synapses expressing slow AMPA (Pampaloni et al., 2021). Such detonating synapses have been suggested to be involved in memory formation (McNaughton and Morris, 1987) and are actively explored in the hippocampus (Vargas-Barroso et al., 2026). Perhaps the most interesting aspect of our results is that CNIH3 overexpression in dCA1 reveals a hidden mosaic of postsynaptic sites, some of which accept and some which reject slow AMPARs. The physiological meaning of this “silent” diversity is currently unclear.

The role of CNIH3 on AMPA kinetics has been tested in GluA2-containing complexes (Schwenk et al., 2009; Shi et al., 2010; Coombs et al., 2012; Herring et al., 2013; Shanks et al., 2014; Brown et al., 2018; Hawken et al., 2017). Here, by co-expressing the CNIH3 with a variety of GluA subunits in HEK293T cells, we also report how the CNIH3 is powerfully slowing both homomeric and heteromeric receptor kinetics (Fig. 4), to a greater extent than almost any other auxiliary subunit. Interestingly, unlike (Coombs et al., 2012), who report a stronger effect in GluA2(Q)-containing receptors, we saw slowing of decay and block of desensitization independent of the GluA subunit. Overall, CNIH proteins do not present a unique electrophysiological signature by which their presence can be inferred. CNIH proteins (CNIH2/3) have been proposed to selectively affect GluA1-containing receptors in hippocampal slices, as the KD of CNIH2 has no effect on GluA1 KO hippocampal slices (Herring et al., 2013). Nonetheless, the specific role of CNIH3 on AMPAR kinetics in its native environment is understudied, usually proposed to be redundant with CNIH2, and only a GluA trafficking factor (Herring et al., 2013). For instance the double KO of CNIH2 and CNIH3 shows a slightly worsened phenotype compared to the single KO of CNIH2, a midly faster decay of mEPSC in mouse hippocampal slices (Herring et al., 2013). Notably, in currently available structures of AMPA receptors derived from the brain, it is not possible to distinguish between CNIH2 and CNIH3 (Yu et al., 2021). Perhaps masking an important sub-population, this level of regulation might be invisible to connectomics approaches, despite being highly decisive for synaptic strength. Other previous work on CNIH could have trivially missed the effect of KD by using dorsal regions. Recent structural studies show that endogenous CNIH-family members are associated well enough to AMPA receptors in HEK-cell expression to be trapped in AMPA receptor complexes resolved by cryo-EM (Yen et al., 2026), perhaps explaining some of the previous variable results in HEK cells.

In CA1 pyramidal neurons, application of glutamate can also drive the activation of other conductances. Importantly, kainate receptors are expressed in CA1 pyramidal neurons and respond to a kainate application (Bureau et al., 1999), a response which is absent in the GluK2 -/- mice. These receptors display slower kinetics than AMPAR, such as in CA3 pyramidal neurons resembling the properties of the current we described in this work (T_decay_ > 100ms, (Cossart et al., 2002; Castillo et al., 1997). The GluK1-specific blocker UBP 310 that we use rule-out the involvement of GluK1 homomer, but GluK2-containing receptors could be incompletely blocked (Pollok and Reiner, 2020). Nonetheless, to our knowledge, synaptic kainate receptors were not reported in CA1 pyramidal neurons (Evans et al., 2019; Cossart et al., 2002). Therefore, the involvement in the slow current we recorded in CA1 during SC stimulation, correlating what we described with MNI-glutamate uncaging (Fig. 2), is unlikely to involve kainate receptors. Moreover, we previously showed that GYKI 52466, AMPAR-specific blocker, can almost completely abolish these currents (also described in Fig. 2 and Fig. 7), which was not the case in CA3 pyramidal neurons (Pampaloni et al., 2021). Finally, currents in CA1 pyramidal neurons are regulated by AMPAR auxiliary subunits (γ-8 in our previous work, CNIH3 in this study). Altogether these observations show that the involvement of kainate receptors in the slow AMPA current described here, is extremely unlikely.

The apparent strengthening of excitatory transmission by CNIH3 that we report can be expected to have wide consequences. CNIH3 is expressed in motor cortex (Bakken et al. 2021) and was proposed to be involved in different neuropathologies, including opioid dependence (Nelson et al., 2016; Lintz et al., 2026) and epilepsy (Leu et al., 2025). AMPARs might be associated with epileptogenesis in humans, and CNIH3 is upregulated (in common with a number of AMPAR subunits) in principal cells and interneurons of the temporal cortex L5/L6 of epileptic patients (Pfisterer et al., 2020). However, the underlying mechanisms of CNIH3 in excitatory transmission were until now unclear. Recent work reveals a role of CNIH3 in spatial memory of rodents (Frye et al., 2021; Lintz et al., 2026). The overexpression of CNIH3 in the dorsal hippocampus improved spatial memory while the Cnih3 –/–mice displayed a reduced short-term spatial memory. We show that the CNIH3 overexpression increases the slow AMPA distribution, and that the KD of CNIH3 can reduce the abundance of slow AMPA. Therefore a dysregulation of the slow AMPA in CA1 is a good candidate to explain the reported results linking CNIH3 and spatial memory in rodents. Moreover, the lack of effect of CNIH3 KO reported in dCA1 neurons mEPSC (Frye et al., 2021), is consistent with our observation that the CNIH3-induced slow AMPAR kinetics is confined in vCA1 and barely present in dorsal CA1. The same authors did not observe any change in AMPAR subunits in the ventral hippocampal synaptosomal fraction from Cnih3 –/– mice, or any variation of CNIH2 expression, suggesting that our KD approach would probably target CNIH3-AMPA complexes without affecting the synaptic GluA subunits composition or CNIH2 in the hippocampus.

Our work demonstrates one potential reason as to why the slow AMPA current has evaded popular consciousness for so long. If studies were confined to the dorsal region of the hippocampus, slow AMPA is so rare that one would never credit it as a real phenomenon. However, slow AMPA currents appear frequently in other brain regions, including striatum (Lape and Dani, 2004), cerebellar Purkinje cells (Devi et al., 2016), unipolar brush cells (Lu et al., 2017) and more (for a review see (Pampaloni and Plested, 2022). It is currently unclear whether selective CNIH-family expression is responsible, or if other subunits or factors contribute to even wider AMPAR synaptic diversity. The selective forebrain expression of γ-8 was already exploited to target a subpopulation of AMPA receptors for therapeutic use (Kato et al., 2016). The distribution of slow AMPA may likewise also have cellular, regional, and subcellular gradients, with implications for brain function and pathological states.

### Limitations of the study

In this work we only assessed the dorso-ventral axis, and did not examine diversity across other axes, such as superficial and deep cells in the stratum pyramidale (Valero et al., 2015). We did not examine ternary combinations of GluA subunits with CNIH3 and TARPs, mainly because the expression of CNIH3 is so toxic for HEK cells, and this situation is exacerbated by TARP coexpression. Further developments in cell culture techniques could be used to address this point. Alternatively, dorsal CA1 is “slow AMPA-free” so mechanisms could be dissected there through overexpression of CNIH3 mutants. Moreover, we have not tested a panel of AMPA receptor subunits to see what other subunits are involved. In particular, we have not explored the role of the Cornichon homolog protein 2 (CNIH2), because it does not display a gradient of expression in CA1, nor a difference of expression between DG and CA1 (Fig. 3). In the hilar mossy cells, where CNIH2 influences the decay, the changes time constant for EPSCs were not so great (about 2-fold as compared to 20-fold) as we observe here with CNIH3 (Boudkkazi et al., 2014). Finally, we provide only circumstantial evidence (molecular genetics and strong modulation of gating) that CNIH3 participates in synaptic slow AMPA receptors. The actual subcellular distribution of CNIH3 remains to be determined.

## Methods

### Mouse organotypic slice

Hippocampi were sampled from P6 to P8 WT C57Bl/6 mice of either sex and dissected as in (Moutin et al., 2020), taking care to remove any remaining cortex attached to the hippocampus. Dorsal and ventral sides were visually recognized and 350 µm-thick transverse slices were cut using McIlwain Tissue Chopper and separated accordingly. The first few slices of the two poles (ventral or dorsal) were cultured separately. Additional precautions were taken to visually recognize the shape of ventral and dorsal slices (see Fig. 1 graphical representations). Slices were prepared on filter paper according to the interface method (Stoppini et al., 1991; De Simoni and Yu, 2006) and cultured as in (Pampaloni et al., 2021). Briefly, 3 slices per cell insert were grown in 1 mL of MEM-based mouse slice culture medium supplemented with 15% Horse Serum; 1x B27; 25 mM HEPES; 3mM L-Glutamine; 2.8 mM CaCl2; 1.8 mM MgSO4; 0.25 mM Ascorbic Acid and 6.5 g/L D-Glucose. The medium was fully replaced each Monday and Friday, independently of the date of slice preparation. Cultures were grown for 16 to 23 days in controlled humidity at 5% CO_2_ in an incubator at 34°C.

### Virus design and expression

AAV-shRNAs were designed against mouse CNIH3 cDNA (using the siRNA Wizard from InvivoGen) by the Viral Core Facility of Charité, Berlin. Particular attention was taken to verify a lack of cross-reactivity to CNIH2.

shRNA CNIH3_1:

5’GATCCgggtttttacccggagtgtgtTCAAGAGacacactccgggtaaaaacccTTTTTTGGAAAT TAAT 3’

shRNA CNIH3_2:

5’GATCCCatcgcctttgacgagctaagaTCAAGAGtcttagctcgtcaaaggcgatTTTTTTGGAAA TTAAT 3’. The shRNAs-induced mRNA reduction were validated by qPCR, giving about 80% (shRNA CNIH3_1) and about 50% (shRNA CNIH3_2) reduction of CNIH3 mRNAs in primary cortical culture.

AAV2/9 vectors containing shRNA sequences together with a GFP reporter and expressed under the U6 promoter were designed by the Viral Core Facility of Charité, Berlin. For AAV-CNIH3, the mouse CNIH3 cDNA was cloned together with GFP resulting in the following plasmid: pAAV-CAG-mCNIH3-IRES-GFP. For AAV-GFP, the plasmid pAAV-CAG-GFP was used for viral production.

The resulting AAVs were diluted in 15uL of medium and applied at directly on top of a ventral (shRNAs, at DIV1) or dorsal (CNIH3/GFP overexpression, at DIV7) slice in the following quantities: AAV-Scrambled ≈ 2.5e^9^ Viral Genomes (VG), AAV-shRNA-CNIH3_1 ≈ 2.1e^9^ VG, AAV-shRNA-CNIH3_2 ≈ 2.4e^9^ VG, AAV-CNIH3 ≈ 5.5e^8^ VG and AAV-GFP ≈ 2.9e^9^ VG.

All the viruses were stored at –80°C and each aliquot was thawed once and used immediately.

### Whole-cell patch clamp in organotypic slices

Slices were superfused at 2 mL/min at room temperature with recirculating HEPES-based artificial cerebrospinal fluid (aCSF) containing 145 mM NaCl, 2.5 mM KCl, 2 mM CaCl_2_, 1 mM MgCl_2_, 10 mM HEPES and 10 mM glucose, with adjusted osmolarity ≈ 310 mOsm/L and pH ≈ 7.3 with NaOH. 4-methoxy-7-nitroindolinyl-glutamate (MNI-caged-L-Glutamate, 0.5 mM; HelloBio HB0423), and a cocktail of different non-AMPAR blockers were added to the aCSF: AP-5 (HelloBio; 20 µM), SR-99531 (HelloBio; 10 µM), (RS)-MCPG (HelloBio; 200 µM) and UBP 310 (HelloBio; 10 µM). For recordings of Fig. 1 Supp. 3, TTX (HelloBio; 1 µM) was also added. The drugs were perfused at the beginning of the day and used for the whole day of recordings. For GYKI 52466 (HelloBio, 100 µM) experiments, the drug was applied at 1 mM (10x final content) together with 1 mL of aCSF solution directly in the recording chamber, and the subsequent recordings were performed after waiting for a few minutes for the drug diffusion.

Microelectrodes with tip resistances of 3–8 MΩ were pulled from borosilicate glass capillaries (1B150F-4, World Precision Instruments) using a Sutter P-1000 puller, filled with the AlexaFluor-594 dye (Leica Microsystems; 20 µM) together with an intracellular solution containing 135 mM K·CH_3_SO_3_, 4 mM NaCl, 2 mM MgCl_2_, 2 mM Na_2_ATP, 0.3 mM Na_2_GTP, 0.06 mM EGTA, 0.01 mM CaCl_2_, 10 mM HEPES, adjusted to 300 mOsm/L and pH 7.2-3 and kept on ice. For recordings of Fig. 1 Supp. 3, the K·CH_3_SO_3_ was replaced by Cs·CH_3_SO_3_ at the same concentration (135 mM). For the experiments involving stimulation of the Schaffer Collateral, QX-314 (HelloBio, 5 mM) was added to the pipette solution.

Somatic whole-cell patch clamp recordings of visually-identified CA1 pyramidal neurons and dentate gyrus granule cells were performed at 16-23 days of slice culture. The pipette was brought into the slice while delivering a positive pressure of ≈ 50 mPa. Cells directly at the slice surface were generally avoided, as the neuronal structure was affected by the hippocampal cut. Selected cells were therefore at a roughly estimated depth of ≈ 30-150 µm from the surface. Negative pressure was applied to reach the gigaseal (R > 1 GΩ) where pipette capacitance was rectified.

Whole-cell voltage clamp was performed at V_h_ = –60 mV. To monitor the uncompensated series resistance, a depolarizing voltage step (+10 mV, 50 ms) was delivered at the beginning of each sweep of a procedure (uncaging or electric stimulation). Liquid junction potential was not rectified, and recordings with R_access_ > 50 MΩ and leak > 200pA were discarded. All the recordings were done at room temperature (≈ 23 °C).

Data were acquired with a Multiclamp 700B amplifier (Molecular Devices), digitized at 10 kHz by Dendrite (Sutter Instrument) and computed in Igor Pro 9 (Wavemetrics).

### Glutamate uncaging and Schaffer Collateral stimulation

Glutamate was uncaged by a 405 nm laser diode (one-photon excitation) mounted on a custom-built upright microscope (Scientifica SliceScope). The collimated beam was directed through a UGA-42 Firefly laser scanner (Rapp Optoelectronic GmbH), and directed through a Zeiss epifluorescence reflector (Examiner A1) into the water dipping 60x objective (Olympus LUMPlanFL N; N.A. 1) as previously described. The uncaging laser and ORCA-Flash 4.0 (Hamamatsu) camera were controlled by SysCon software (Rapp) and ImageJ (https://imagej.nih.gov/ij/) respectively. After reaching the whole cell modality, we waited a few minutes to allow the diffusion of AlexaFluor-594 dye into the processes of the neuron. Afterward, dendritic regions were selected by illuminating the sample with a 595 nm LED (Thorlabs). UV light pulses of 1ms (10Hz) were delivered by triggering the laser (at 50% power, delivering a 5.73 mW pulse) on the selected location. Simultaneous passage of orange excitation light (600 nm) and 405 nm light for uncaging was achieved by using a 405/488/594nm Laser Triple Band filter set (TRF 69902; Chroma) mounted in a Zeiss TIRF cube. At the end of each experiment, we documented the uncaging sites using a tiled, multifocal plane fluorescence micrograph of the filled neuron.

To evoke whole cell EPSCs in CA1 principal cells from stimulating the Schaffer Collateral, we used an 1-2MΩ electrode when filled with aCSF. The stimulating electrode was placed at the slice surface in the CA3/CA1 interface at the stratum radiatum to activate Schaffer collateral/commissural afferents (150-400 µm away from the recording electrode). Bipolar stimulation was applied with an Iso-Flex constant-current stimulator (API Instruments, Jerusalem, Israel), and the stimulation trigger (10 x 1 ms pulses at 10 Hz) was controlled by Igor software. The power of the stimulator was increased until the EPSCs amplitude recorded in whole-cell patch clamp reached 50-150 pA.

Glutamate uncaging and SC stimulation protocols were delivered 5 times, and the resulting traces were averaged before being low-pass filtered at 0.5 kHz. The voltage steps at the beginning of each stimulation were also delivered 5 times and averaged then low-pass filtered at 1kHz.

### Single-cell RNAseq analysis

For the Fig. 3, the RNAseq data were extracted from the freely available HippoSeq dataset of the Janelia Research Campus (https://hipposeq.janelia.org), displaying the FPKM measurement of excitatory neurons of dorsal and ventral microdissected mouse hippocampi (Cembrowski et al., 2016a). Ratio of Fragments Per Kilobase Millions (FPKM) was done from the mean of three replicates of each selected transcripts. As recommended by (Cembrowski et al., 2016a), only transcripts displaying a substantial abundance (mean FPKM > 10) were selected, and a strict analysis of differences displaying only 3-fold differences were applied. For the Fig. 3 Supplementary Fig. 1, data were extracted from the freely available Smart-Seq v4 and 10x Chromium dataset of the Allen Institute (https://celltypes.brain-map.org/rnaseq/mouse_ctx-hpf_smart-seq?selectedVisualization=Heatmap&colorByFeature=Cell+Type&colorByFeatureValue=Gad1). For further details, including differences in data collection and processing, see (Yao et al., 2021). The Trimmed Mean Expression (TME) of the selected transcripts were sorted and averaged from the following classified clusters: CA1 (clusters 337-345), DG (clusters 361-364), vCA1 (clusters 334-336) and dCA1 (clusters 346-348).

### HEK293 cell culture

Human embryonic kidney (HEK) 293 cells were obtained from the German Collection of Cell culture, cultured with Minimum Essential Media (PAN Biotech GmbH) supplemented with 6% FBS (PAN Biotech GmbH), and maintained at 37°C, 95% air and 5% CO2 in a humidified incubator. Three days prior to transfection, cells were seeded on 10 mm glass cover slips coated in poly-L-lysine (Sigma Aldrich). Cells were transfected with PEI Following standard protocol. A total of 1 μg DNA plasmid was used for all transfections. In cases of co-transfection, equal ratios of all plasmids were used. Receptor subunit plasmids contained the subunit coding sequence and GFP under the IRES sequence. The CNIH3 plasmid contained the auxiliary protein coding sequence in the pRK vector expressing mCherry also following an IRES. After 6 hours the transfection was stopped by removal of transfection media followed by two washes with PBS. Transfected cells were then cultured in MEM without glutamine (Fischer Scientific) containing 6% FBS (PAN Biotech GmbH), GlutaMAX (Thermo Fisher Scientific) and 40 μM NBQX (Sigma Aldrich). Electrophysiology experiments were carried out 24-48 hours later.

### HEK293 electrophysiology

Positively-transfected cells were assessed by their green and red fluorescence as appropriate. Cells were patched in an outside-out configuration using glass pipettes pulled to 4-6 MOhm resistance when filled with internal solution. Extracellular solution contained (in mM): 150 NaCl, 0.1 MgCl_2_, 0.1 CaCl_2_ and 5 HEPES, titrated to pH 7.3 with NaOH, 300 mOsm. Intracellular solution contained (in mM): 115 NaCl, 10 NaF, Na_4_BAPTA, 0.5 CaCl_2_, 1 MgCl_2_, 5 HEPES and 10 Na_2_ATP, titrated to pH 7.3 with NaOH, 290-300 mOsm. Both solutions were filtered through a nylon 0.2 μM filter with intracellular solution aliquots stored at −20°C. Freshly thawed aliquots were subsequently used and kept on ice throughout the patching day. Application of 10 mM glutamate was performed using a fast-perfusion system described previously (Plested and Poulsen, 2021). Briefly, a four-square glass barrel tool was mounted on a piezo stack (Physik Instrument) for brief 1 ms or 500 ms pulses of glutamate. To test inhibition by GYKI 52466, (100 μM), the drug was added to both control and glutamate barrels. Currents were measured at a holding potential of −60 mV for kinetic analysis and +50 mV for GYKI 52466 inhibition (to match our previous study, (Pampaloni et al., 2021). Membrane patch currents were low-pass filtered at 5 kHz using an Axopatch 200B amplifier (Molecular Devices) and acquired with Axograph X software (Axograph Scientific).

### Data Analysis

Recordings and figures were analyzed and made using Igor (WaveMetrics). Steady-state current was calculated as a percent of the peak response. Desensitization and decay was determined by fitting the traces with a sum of two exponentials with constants expressed as weighted mean of multiple components. The resulting T_weighted_ was obtained by computing the A_fast_ and A_slow_ amplitude together with the T_fast_ and T_slow_ kinetics of the double exponential: 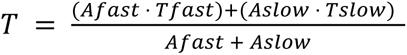. For neuronal recordings, the 5 sweeps were automatically averaged by a custom-built program. Statistical analysis were carried out using Prism 10 (Graphpad). Graphs were made using Igor Pro 9 (WaveMetrics).

## Acknowledgements

We thank the viral core facility of the Charite (Thorsten Trimbuch and colleagues) for the design and production of AAV constructs for shRNA and overexpression. N.P.P. was recipient of an EMBO Long Term Fellowship, J.D.N. was recipient of a NSERC. Canada Fellowship. A.J.R.P. received funding from the Deutsche Forschungsgemeinschaft (DFG, German Research Foundation) under Germany’s Excellence Strategy – EXC 2049 – 390688087 (NeuroCure Cluster of Excellence). We thank James Gowman for providing an Igor Procedure for the automated redimension and averaging of the traces.

## Author contributions

Conceptualization, M.B., J.D.N., N.P.P., and A.J.R.P.; methodology, M.B., J.D.N., N.P.P., and A.J.R.P.; Investigation, M.B., J.D.N., and N.P.P..; writing – original draft, M.B. and A.J.R.P.; writing – review & editing, M.B., J.D.N., N.P.P., and A.J.R.P.; funding acquisition, J.D.N., N.P.P., and A.J.R.P.; resources and supervision, A.J.R.P.

## DECLARATION OF GENERATIVE AI AND AI-ASSISTED TECHNOLOGIES IN THE WRITING PROCESS

We did not use generative AI or AI-assisted technologies during the preparation of this work.

