## Supplementary Figures for "CNIH3 is a molecular signature of slow AMPA receptors"

### Supplementary data:

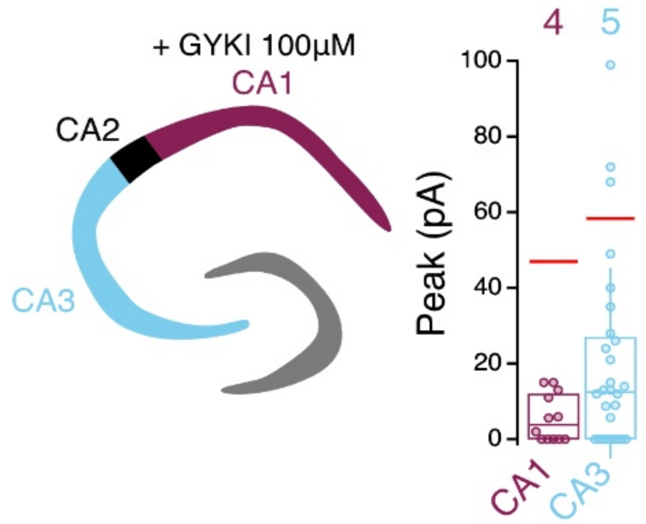


**Figure 1. Supplementary Fig. 1 Incomplete GYKI 52466 inhibition of MNI-glutamate uncaging responses in CA3 of the ventral hippocampus**

After incubation with AMPAR selective blocker GYKI 52466 (100µM), the peak of the response to a glutamate uncaging (1 ms UV pulse, 5.73 mW), were recorded in pyramidal neurons of CA1 and CA3 in ventral hippocampal organotypic slices. Responses in CA1 were almost non-existent (5.6 ± 1.8pA, 88% reduction compared to the mean measured in other slices without GYKI, n = 12 sites, 4 cells). However, in CA3, responses were on average substantially larger (20.2 ± 4.8 pA, 60% reduction compared to CA3 without GYKI, *n* = 28 sites in 5 cells). Further, substantial responses >50 pA were readily detected, probably corresponding to Kainate receptors that were spared by UBP 310. The nature of the stimulation (mono or poly-dendritic) in GYKI 52466-incubated slices was not monitored, so these results are an approximation and the inhibition could be even stronger (see (Pampaloni et al., 2021) for CA1). In some but not all cases, the responses of CA1 and CA3 pyramidal cells in GYKI were recorded from the same slice, therefore confirming both the complete action of GYKI in CA1, and a population of GYKI- and UBP310-resistant receptors in CA3. These responses were almost universally “slow” by our decay >25 ms, >10% steady-state criteria.

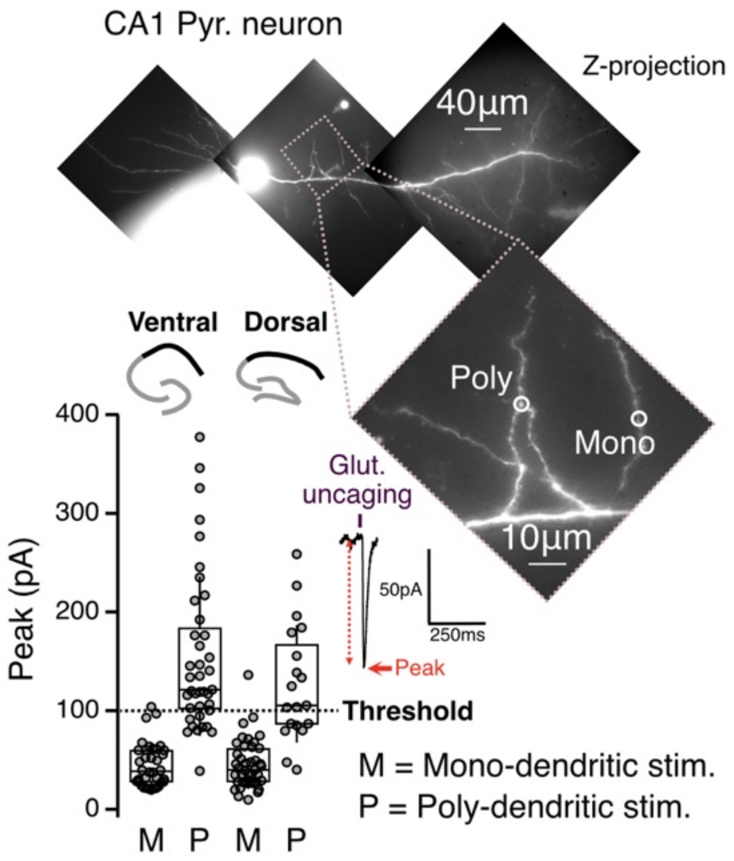


**Figure 1. Supplementary Fig. 2 Poly-dendritic responses can be discarded with a threshold**

The peak of the response to glutamate uncaging (1 ms 405 nm laser exposure, 5.73 mW under the 60x objective) adjacent to identified dendrites in hippocampal organotypic slices and recorded in whole-cell patch clamp displayed a range of amplitudes. Responses from ventral and dorsal CA1 were compared. As discussed in the text, one-photon uncaging is relatively unprecise in the z-axis. Using the z-projection of tiled fluorescence micrograph of the recorded neurons, the stimulations were divided into two categories: mono-dendritic (M, laser targeting only one dendrite in the z-stack), and poly-dendritic (P, laser spot adjacent to one dendrite or more from the same cell, located in different z-planes). The peak of mono-dendritic responses in ventral CA1 (44.9 ± 3.73 pA, $n$ = 35 sites) showed significantly lower amplitude compared to the peak of presumed poly-dendritic responses (155.2 ± 12.9 pA, $n$ = 38 sites), $p<0.0001$ (unpaired t-test). Likewise, the peak of mono-dendritic responses in dorsal CA1 (45.6 ± 3.9 pA, n = 40 sites), showed significantly lower amplitude compared to the peak of poly-dendritic responses (127 ± 13.7 pA, n = 18 sites), $p<0.0001$ (unpaired t-test). By thresholding the responses at 100 pA, we could effectively select mono-dendritic responses.

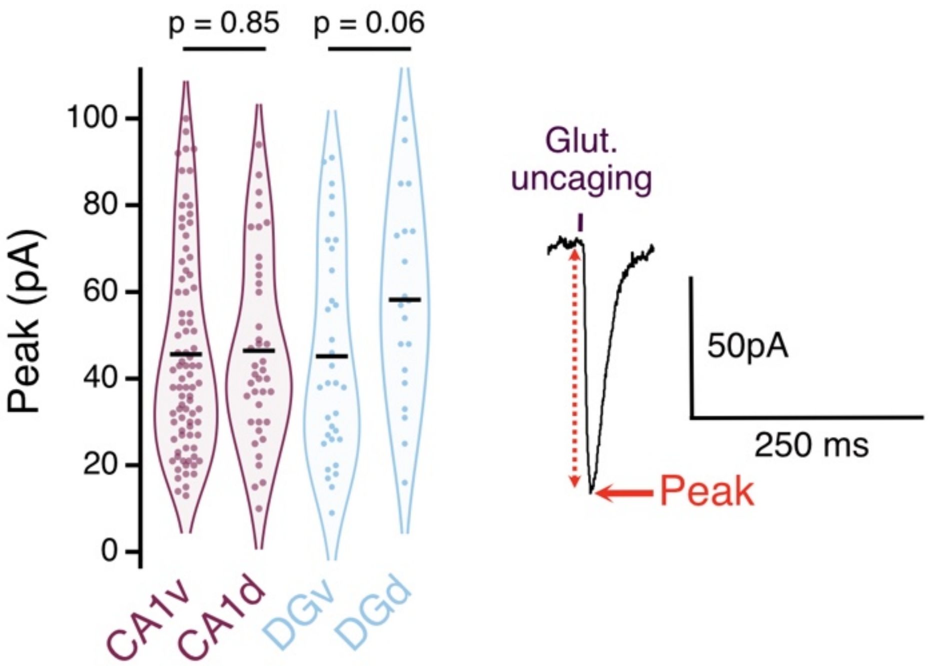


**Fig. 1. Supplementary Fig. 3 Peak of responses in pyramidal neurons of CA1 and granule cells of Dentate Gyrus**

The peak of the response to a 1 ms UV pulse (5.73 mW) uncaging glutamate were recorded in CA1 pyramidal cells (ventral and dorsal, purple) and Dentate Gyrus (DG) granule cells (ventral and dorsal, blue) using whole-cell patch clamp. Responses greater than 100 pA were discarded as being probably from multiple dendrites (see Fig. 1 Supp. Fig. 2). Peak amplitudes showed no appreciable difference between ventral and dorsal hippocampal CA1 pyramidal cells (ventral: 45.6 ± 2.5 pA; dorsal: 46.4 ± 3.4 pA; unpaired t-test) or Dentate Gyrus (ventral: 45.15 ± 4.2 pA; dorsal: 58.15 ± 5.2 pA; unpaired t-test).

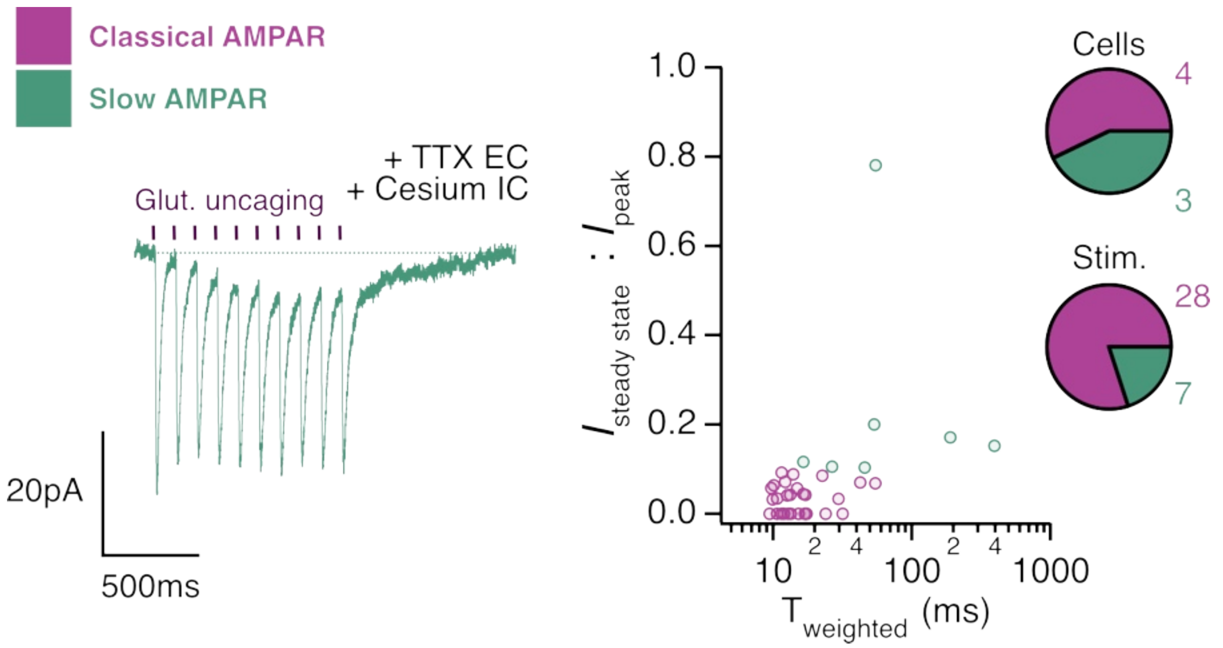


**Figure 1. Supplementary Fig. 4 $Na_v$ and $K^+$ channels are not required for slow AMPA currents in ventral CA1**

The hippocampal organotypic slices were incubated with 1 µM of TTX, blocking $Na_v$ channels, and the intracellular medium were switched to a Caesium-based solution, blocking $K^+$ channels. Slow responses were recorded with glutamate uncaging (ten UV 1ms pulse 5.73mW at 10 Hz) as displayed on the left, with 25% of slow responses ($I_{ss}$ : $I_{peak}$ > 0.1 and $T_{weighted}$ > 25ms) . green: slow responses; purple: classical responses. *n* = 35 stim., 7 cells, 7 slices.

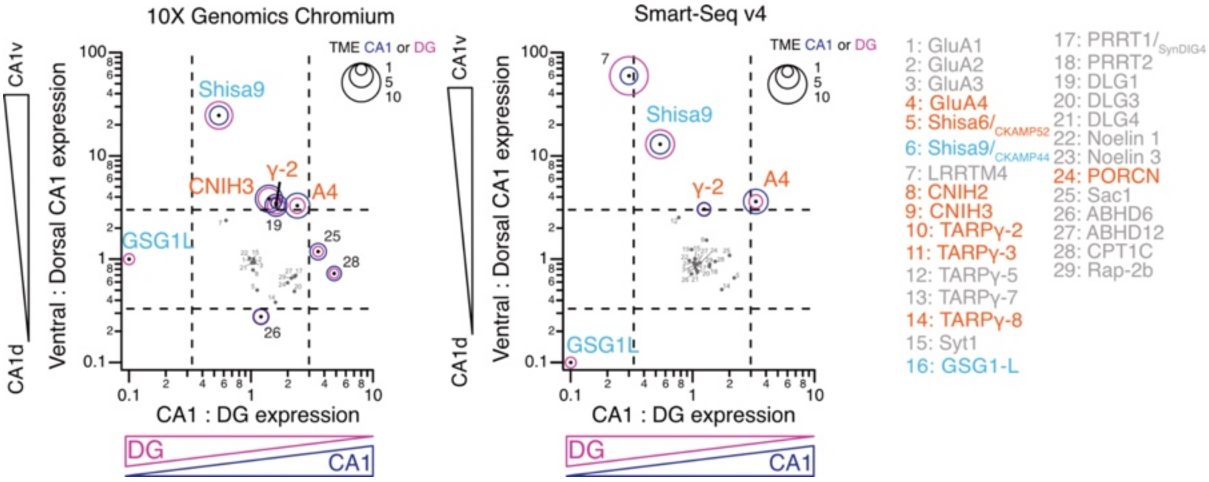


**Figure 3. Supplementary Fig. 1 Analysis of AMPAR components in two additional transcriptomics datasets**

Single-cell RNA expression of glutamatergic neurons of the adult mice brain from a taxonomy of transcriptomic cell types across hippocampal formation (https://celltypes.brain-map.org/rnaseq/mouse_ctx-hpf_smart-seq?selectedVisualization=Heatmap&colorByFeature=Cell+Type&colorByFeatureValue=Gad1 Yao et al. 2021). The selected transcripts are the same as Fig. 3, and the corresponding number in the graph is listed on the side. The abundance of each transcript reported in Trimmed Mean Expression indicated by circles in CA1 (blue) and DG (purple). The ratio of the TMEs of Ventral and Dorsal CA1 (Y axis) is plotted together with the ratio of the TMEs of CA1 and DG (X axis). The dotted lines indicate the threshold upon which a difference is considered as meaningful (more than 3- fold enriched or depleted). The known positive (orange) and negative (blue) regulators of AMPA decay kinetic relevant for this study are indicated (orange: slowing decay, blue: speeding decay). CNIH3 is a hit in 10x Chromium dataset (as in HippoSeq data, Fig 3) but does not present a dorsal:ventral gradient in the Smart-Seq v4 data, perhaps reflecting the different ways in which these data were collected and processed.

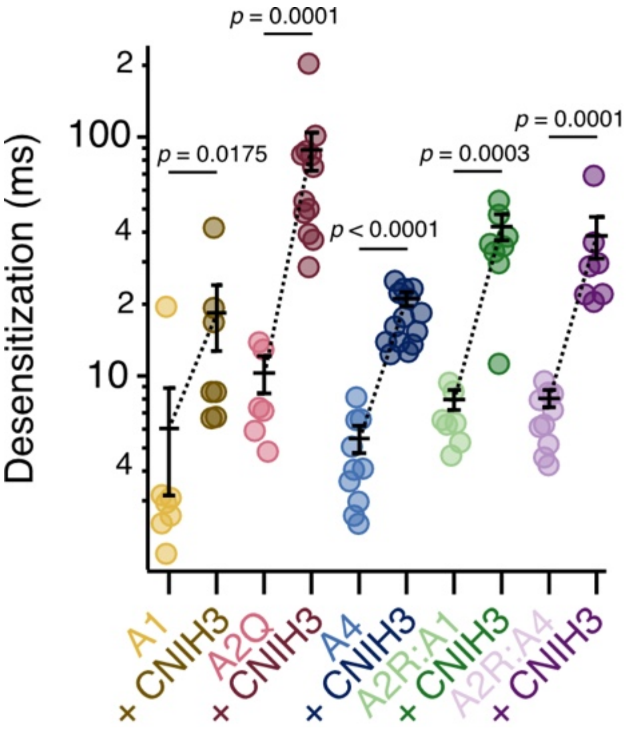


**Figure 4. Supplementary Figure 1. Desensitization of AMPAR-CNIH3 complexes**

Effect of CNIH3 on AMPAR desensitization during a long (500 ms) glutamate exposure. Desensitization is significantly slowed for all homomeric and heteromeric AMPARs in the presence of CNIH3; GluA1 (from 6.0 ± 2.9% to 18.4 ± 5.7%, $p = 0.0175$), GluA2Q (from 10.3 ± 1.8% to 88.5 ± 16.1%, $p = 0.0001$), GluA4 (from 5.5 ± 0.7% to 21.1 ± 1.4%, $p < 0.0001$), GluA2R:1 (from 8 ± 0.8% to 42.3 ± 5.0%, $p = 0.0003$), and GluA2R:4 (from 8.1 ± 0.7% to 37.7 ± 7.3%, $p = 0.0001$), without and with CNIH3, respectively. Representative recordings shown in Figure 4. Error bars represent standard error of the mean. Statistical significance was evaluated using the Mann-Whitney test.

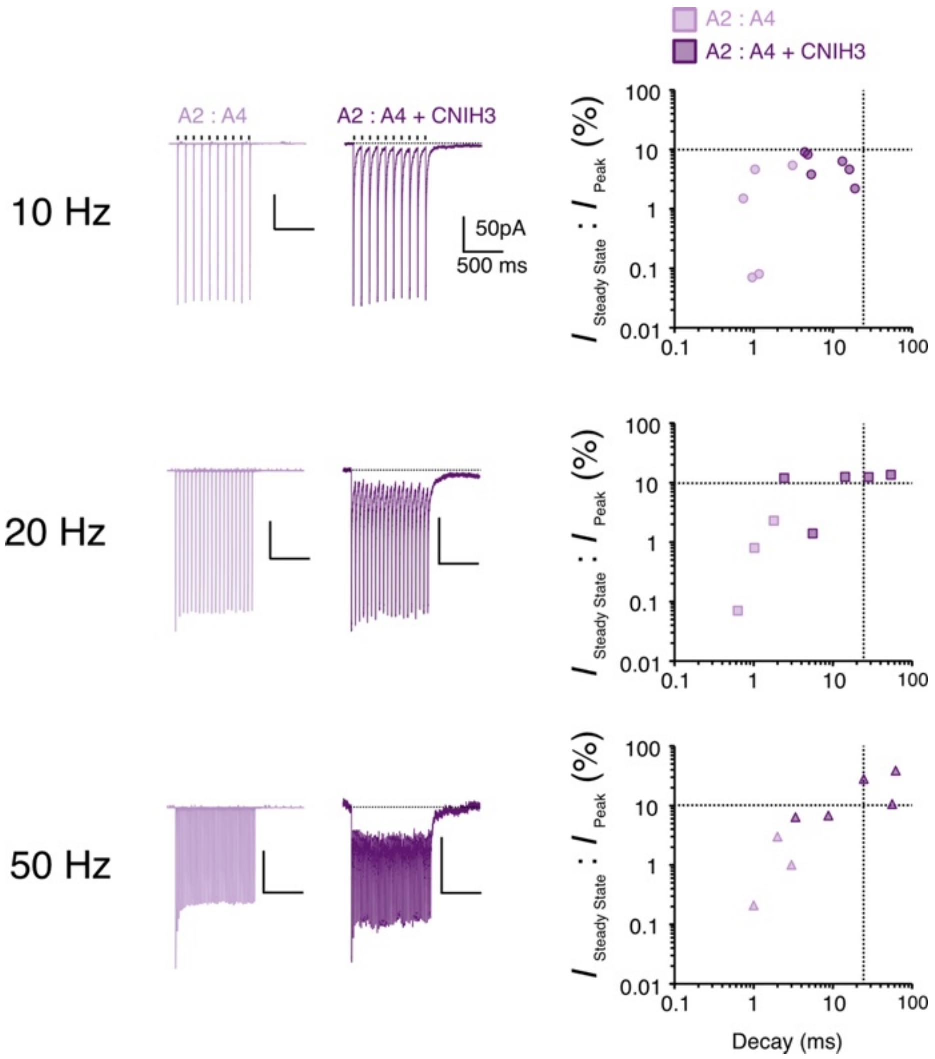


**Figure 4. Supplementary Figure 2. Trains of glutamate stimulation can partially mimic the slow responses described in neurons.**

Representative recordings of current responses from a heteromeric AMPAR (GluA2R:4) during train pulses of 1 ms glutamate at different frequencies. Two-dimensional plots with $I_{ss}$ normalized to the 1$^{st}$ peak response ($I_{steady\ state}$ : $I_{peak}$) plotted with the associated decay ($T_{weighted}$ of the last response). At all frequencies, CNIH3 co-expression slowed AMPA receptor kinetics. *Left*: Receptor alone, *Right*: Receptor co-expressed with CNIH3. At low frequency (10 Hz, *Top*) the slow kinetics of receptors with CNIH3 begin to become apparent. At higher frequencies (20 Hz, *Middle*) the high steady state and slow decay is indicated which is more pronounced at 50 Hz (*Bottom*).

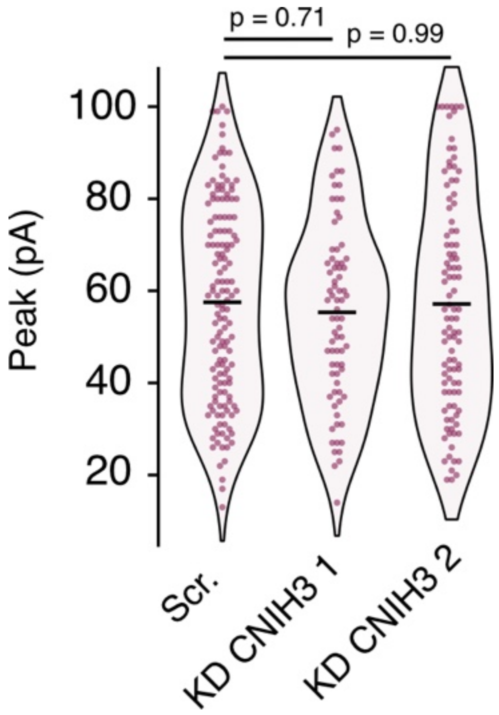


**Figure 5. Supplementary Fig. 1 Effect of shRNAs on the peak of responses in ventral CA1**

The peak of the response to a 1ms UV pulse (5.73 mW) uncaging glutamate was recorded in CA1 pyramidal cells using whole-cell patch clamp. Responses greater than 100 pA were discarded as being probably from multiple dendrites (see Fig. 1 Supp. 2). Peak amplitudes showed no appreciable difference between the different transduced shRNA (Scr: 57 ± 1.8pA; shRNA 1: 55 ± 2.2pA and shRNA 2: 57 ± 2.3pA). Statistic significance was tested using Dunnett's multiple comparisons test with adjusted p-value indicated on the graph.

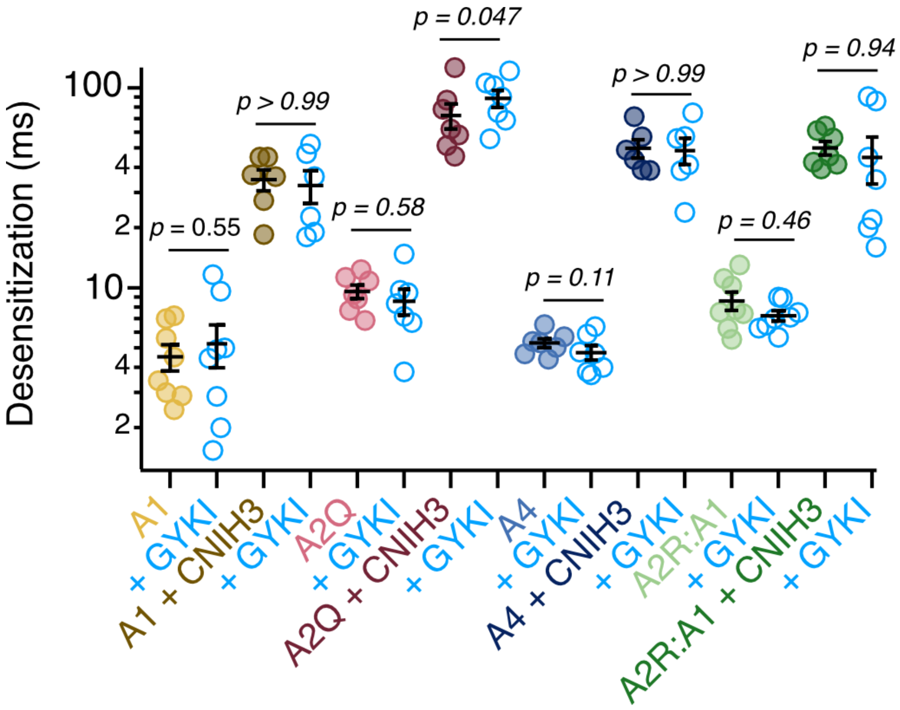


**Figure 7. Supplementary Figure 1. Effect of GYKI 52466 on the desensitization of AMPAR-CNIH3 complexes**

Effect of GYKI 52466 on AMPA receptor desensitization, with and without CNIH3 coexpression. In nearly all AMPA receptors, the receptor desensitization is not altered with GYKI 52466 application, addition of CNIH3 does not alter the desensitisation with GYKI 52466; GluA1 (from 4.5 ± 0.7% to 5.3 ± 1.3%, p = 0.55), GluA1 + CNIH3 (from 34.8 ± 4.2% to 32.5 ± 6.1%, p >0.99), GluA2Q (from 9.6 ± 0.8% to 8.57 ± 1.3%, p = 0.58), GluA4 (from 5.3 ± 0.3% to 4.8 ± 0.4%, p > 0.11), GluA4 + CNIH3 (from 49.9 ± 5.2% to 48.6 ± 7.3%, p > 0.99), GluA2R:1 (from 8.6 ± 0.9% to 7.3 ± 0.4%, p = 0.46), GluA2R:1 + CNIH3 (from 50.1 ± 3.9% to 44.9 ± 11.8%, p = 0.94), with glutamate and with glutamate + GYKI 52466, respectively. For GluA2Q + CNIH3, desensitization was slightly slower with GYKI 52466;(from 72.7 ± 10.5% with glutamate to 88.6 ± 8.8%, p = 0.047 with glutamate + GYKI 52466. Error bars represent standard error of the mean. All the statistics were done using the Wilcoxon matched-pairs signed rank test.
